# A molecular circuit integrates environmental cues to coordinate the exit from developmental dormancy

**DOI:** 10.64898/2026.10.24.760914

**Authors:** Omer Ben Dor, Itai Rozenberg, Tehila Atlan, Erin M. Davis, Marva Bergman, Adi Oron-Gottesman, Tommer Argaman, Walter Sandt, David H. Meyer, Yoav Rapoport, Mazal Tawil, Björn Schumacher, Jason E. Podrabsky, Michal Rabani, Itamar Harel

## Abstract

Diapause is a reversible dormancy state that enables organisms to survive extreme environments. In annual killifishes, embryos use diapause to bridge seasonal drought, but how environmental cues are integrated to trigger exit remains unclear. Here, we combine model-based single-embryo transcriptomics with cross-clade small-molecule screening to define the molecular logic of diapause exit. In the African turquoise killifish (*Nothobranchius furzeri*), high-resolution profiling uncovered a temporal transcriptional cascade, while transcriptomic-clock analysis revealed reduced transcriptional noise during developmental reactivation. Transcriptome-guided drug-enrichment analysis nominated overlapping compounds across two annual killifish clades, leading to a targeted chemical screen that identified shared activators and inhibitors of diapause exit. Functional assays support a model in which HSP90 integrates temperature-dependent cues with steroid-hormone signaling. Together, these findings reveal shared molecular machinery that couples environmental sensing to developmental timing, aligns diapause exit with favorable conditions, and provides a powerful comparative framework for dissecting vertebrate dormancy control.

## Introduction

While most killifishes inhabit permanent streams and lakes, some species, termed “annuals,” occupy seasonal ponds that form during the brief hot–wet season characteristic of tropical and subtropical regions, and disappear during the cold–dry season (**Figure 1a**). To survive this habitat, annual killifishes have evolved a unique life cycle that includes rapid growth, early sexual maturation^1,2^, a compressed lifespan^3^, and the ability to pause embryonic development at up to three distinct stages of embryonic diapause (termed diapause I, II, and III)^4^.

**Figure 1.**
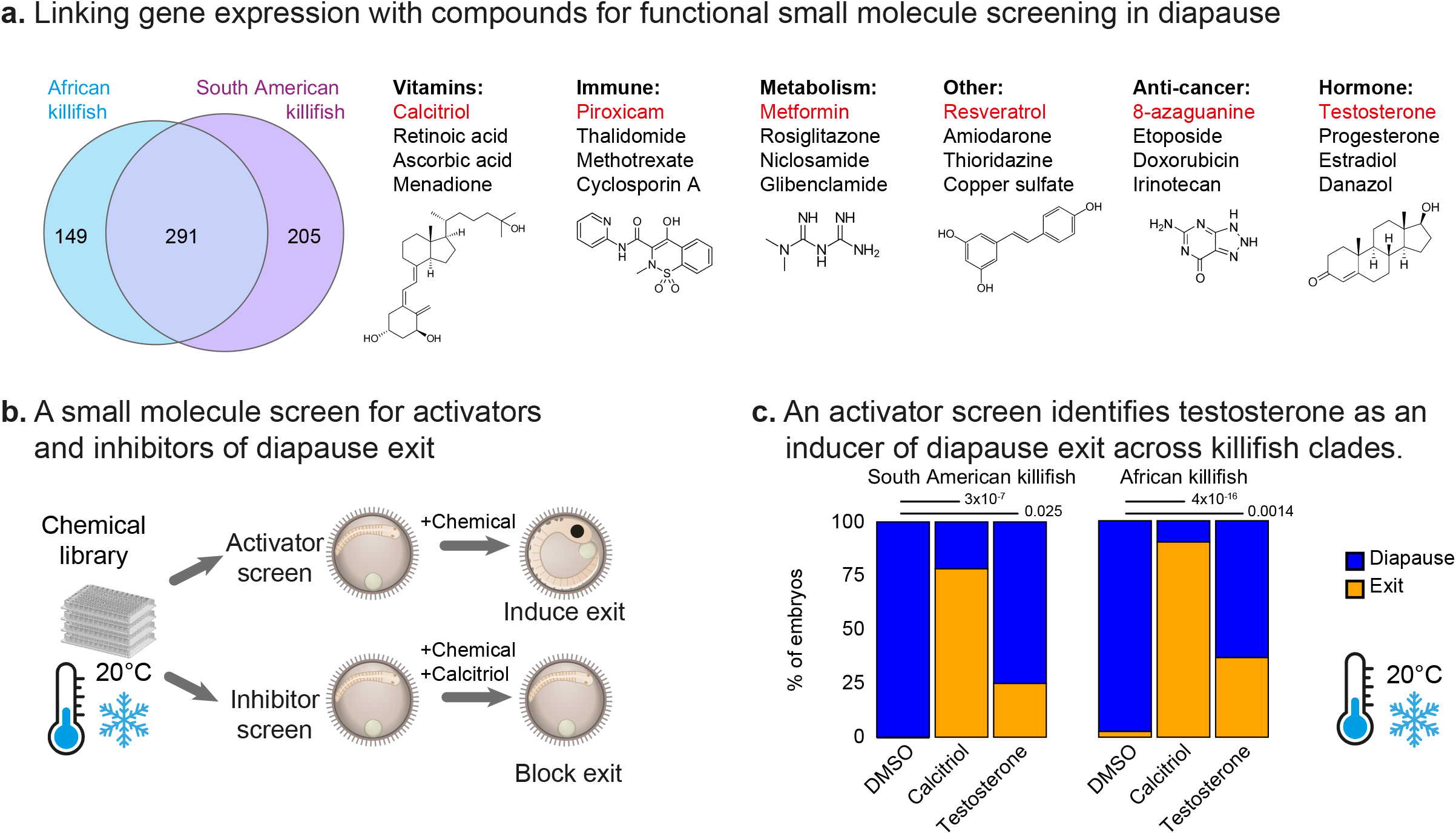
Single-embryo transcriptomic characterization of diapause exit at high temporal resolution in the African killifish. **a.** Schematic of alternative developmental trajectories in the African killifish in relation to primary seasonal cues (precipitation and temperature, based on refs.^3,89^). Three key processes are illustrated: (1) Diapause entry: direct development into diapause; (2) Escape from diapause: bypassing the diapause checkpoint and continuing development; and (3) Diapause exit: resuming development following diapause. The direct-development (or escape) trajectory is well documented under laboratory conditions, whereas its frequency and seasonal timing in natural populations remain uncertain. The diapause trajectory shown represents the predominant annual cycle supported by field studies of wild embryo banks^11^. **b.** A phylogenetic tree of killifish species (based on ref.^22^). Red branches indicate clades that are capable of an annual life cycle with the ability to enter diapause. Numbers indicate millions of years (MYA) since divergence from a common ancestor. **c.** Quantification of cell-cycle re-entry following diapause exit, measured using the FUCCI cell-cycle transgenic reporter. Top: representative images (after ∼5h, 15h, 20h, 25h, 30h post calcitriol exposure). Centre: cell-cycle entry was estimated from GFP signal (y-axis) intensity over a 36-hour period (x-axis), following diapause exit induced by exposure to calcitriol at 20°C. Thin lines represent individual embryos (n = 8), the thick line represents the mean, and the shaded area indicates the standard deviation. Bottom: color bar schematically represents embryo coloration corresponding to RFP/GFP ratios. **d.** PCA of single-embryo RNA-seq data (x-axis: PC2, y-axis PC3). Each point represents one embryo (n = 22); PCA used 10,613 expressed genes. Embryos were collected at 2.5-hour intervals after calcitriol treatment, over a total duration of ∼60 hours. **e.** Heatmap showing log₂ fold changes in gene (rows), temporal expression (columns), relative to time 0 (blue: decrease relative to time 0, red: increase relative to time 0). The color bar below indicates GFP intensity over time (reproduced from panel c) and aligned with the samples’ pseudo-time–corrected order.

Embryonic diapause is a widespread survival strategy^4,5^ used across the tree of life to endure unfavorable conditions^6–9^. In annual killifishes, the major diapause stage used to survive the dry season is diapause II^10,11^. Importantly, embryos can develop along two alternative trajectories to either enter or escape diapause II (**Figure 1a**). In one trajectory, embryos “skip” diapause II (termed “escape” embryos^12^) and develop continuously to the pre-hatching stage. But in another trajectory, embryos can enter diapause II to endure pond desiccation, and then later exit diapause II to resume development when conditions indicate that flooding will again occur.

Mounting experimental evidence supports the view that the escape trajectory is a separate developmental program from diapause II, and not simply faster progression through the same pathway. In particular, differences in heart rate, head shape, optic-cup size, timing of red blood cell differentiation, IGF-I dynamics, and glycolytic and mitochondrial enzyme activities^13–16^ support this view of two separate developmental programs.

Although pond drying and flooding can occur within a rainy season or between rainy seasons, the escape trajectory is primarily documented under laboratory conditions. So far, field surveys of natural *Nothobranchius* dry season embryo banks support a predominantly diapause II state^11^, with development distributed across two seasonal cycles. Selecting between these two trajectories is essential for survival and requires embryos to integrate environmental cues into internal signaling pathways, which ultimately control two key decisions: first, whether to enter or escape diapause II, and second, if and when to exit it. However, which environmental cues are used and how they are integrated into the molecular machinery remain unresolved.

Decisions regarding entry, exit, and escape from diapause II (hereafter diapause) depend on both environmental cues and steroid hormone pathways. Embryos typically resume development before their ponds are reflooded^11^, indicating that diapause exit is triggered by anticipatory cues rather than the flooding of the pond itself. Temperature has also emerged as a key environmental cue in this decision^16,17^. Warmer conditions both favor the escape trajectory and trigger exit from diapause. In contrast, cooler conditions favor the diapause trajectory and promote its maintenance^11,16,17^. Recent work in the South American annual killifish *Austrofundulus limnaeus* further showed that vitamin D_3_ signaling can regulate the choice between diapause entry and escape^17^. Interestingly, vitamin D_3_ has also been linked to other pace-of-life events, including the timing of puberty and the rate of aging traits in mice and humans^18–20^.

Compared to entry and escape from diapause^17,21^, the mechanisms driving diapause exit (**Figure 1a**) remain poorly understood. In particular, the systems-level molecular regulation governing this reactivation remains largely unknown. What specific mechanisms integrate environmental and hormonal inputs to drive the exit process? Furthermore, do similar signals dictate the initial commitment to enter or escape diapause, and why does this trajectory vary across individual embryos^5^? In light of current evidence, steroid hormone signaling is well-positioned to couple environmental state to developmental programs, suggesting a tractable mechanistic axis that may also regulate diapause exit.

Here, we combine high-resolution single-embryo transcriptomics with temporal modeling and chemical perturbation to define the regulatory logic of diapause exit. We identify key transcriptional modules of the diapause-exit trajectory, suggest that these are shared with other clades and with the diapause escape trajectory, and identify an HSP90-dependent mechanism that links temperature and nuclear receptor signaling to developmental reactivation during diapause exit.

## Results

### Vitamin D_3_ promotes both escape and exit trajectories across killifish clades

Exposure of the South American annual killifish *A. limnaeus* to calcitriol, the active form of vitamin D_3_, has been shown to promote the diapause escape trajectory, allowing embryos to develop continuously to the pre-hatching stage^17^. While the African turquoise killifish *N. furzeri,* a distantly related species that shared a common ancestor with *A. limnaeus* approximately 72 million years ago^22,23^ (**Figure 1b**), is known to develop along the diapause and escape trajectories based on temperature^23^, the role of vitamin D_3_ signaling in this response remains unknown.

First, we confirmed calcitriol’s conserved ability to promote the escape trajectory in *N. furzeri* by exposing one-week-old embryos to calcitriol. Whereas 30% (n=60) of control embryos remained in diapause at 28°C, all calcitriol-treated embryos (n=30) followed the escape trajectory. Next, we examined calcitriol’s ability to induce exit from diapause in *N. furzeri* using a FUCCI (Fluorescence Ubiquitination-based Cell Cycle Indicator) reporter line^24^. We first incubated embryos for 3–4 weeks at low temperature (20°C)^17,25^ to synchronize them in diapause II, confirming they arrested in G1 phase of the cell cycle (red fluorescence, **Figure 1c**). Upon calcitriol exposure, we observed a burst of proliferation (green fluorescence, **Figure 1c**), indicating that embryos exited diapause and resumed normal development (**supplementary Movie 1**). Subsequently, >90% of embryos (n=87/96) exited diapause in response to calcitriol. Notably, embryos following the escape trajectory did not show a green proliferation burst, whereas embryos that had briefly entered diapause did (a rapid “in-and-out” treatment induced by a temperature shift, see **Figure S1a**).

Interestingly, time-lapse imaging of FUCCI embryos revealed variability in green fluorescence intensity: fluorescence appeared after 10–12 hours and peaked at 18–22 hours (**Figure 1c**, top, and **supplementary Movie 1**). These data demonstrate that vitamin D_3_ signaling can promote both escape and exit from diapause. However, despite a uniform calcitriol stimulus, exit rates were not fully synchronized across embryos (**Figure 1c**). Instead, analyzing the cohort collectively revealed a several-hour window of embryo-to-embryo variability, highlighting individual differences in the temporal progression of developmental reactivation.

### Model-based single-embryo transcriptomics resolves diapause exit dynamics

We next characterized the transcriptional landscape of vitamin D_3_-induced diapause exit. Diapausing embryos at 20°C were exposed to calcitriol and sampled individually by optimizing RNA extraction from the embryos’ high polysaccharide content^17^ (see **Methods**). Single-embryo samples were collected at high temporal resolution every 2.5 h across ∼60 h of calcitriol-induced diapause exit. RNA-seq profiling recovered 10,613 expressed genes across the entire time-course (**Figures 1d,e**).

Profiling individual embryos, rather than pooled samples, allowed us to capture and partially account for embryo-to-embryo variability in the timing of calcitriol-induced diapause exit observed by FUCCI (**Figure 1c**). Indeed, pseudotime ordering^26^ recovered a coherent developmental progression in PCA space on PCs 2 and 3. This analysis^26^, however, also exposed some inconsistencies, where the transcriptomes of specific samples did not fully match the temporal order in which they were collected (**Figure 1d**). PC1 was largely driven by broad developmental processes such as cell cycle and proliferation (**Figure S1b**), without a meaningful temporal order.

To fully exploit the depth and temporal resolution of our single-embryo dataset, we applied a model-based temporal framework^27^ tailored to analyze densely sampled time-series data^28–31^. We used a hierarchy of three nested parametric functions (**Figure 2a**, **Methods**): constant (null), sigmoid (a single sustained change), and “impulse” (a transient change), which was shown to successfully capture transcriptional dynamics^32^. For each gene, we selected the best-fitting function by using a likelihood ratio test^33^. Genes that reject the constant null hypothesis represent transcripts with temporal changes during diapause exit. We validated that predicted profiles (**Table S1**) closely matched measured expression (98.7% of genes with ‘goodness-of-fit’ test p<0.05, 64.9% with r^2^>0.7), confirming the suitability of this modelling strategy to our data. As an additional validation, we also confirmed its consistency with classical differential expression analysis of grouped time windows throughout the time-course (**Figure S2a-b**).

**Figure 2.**
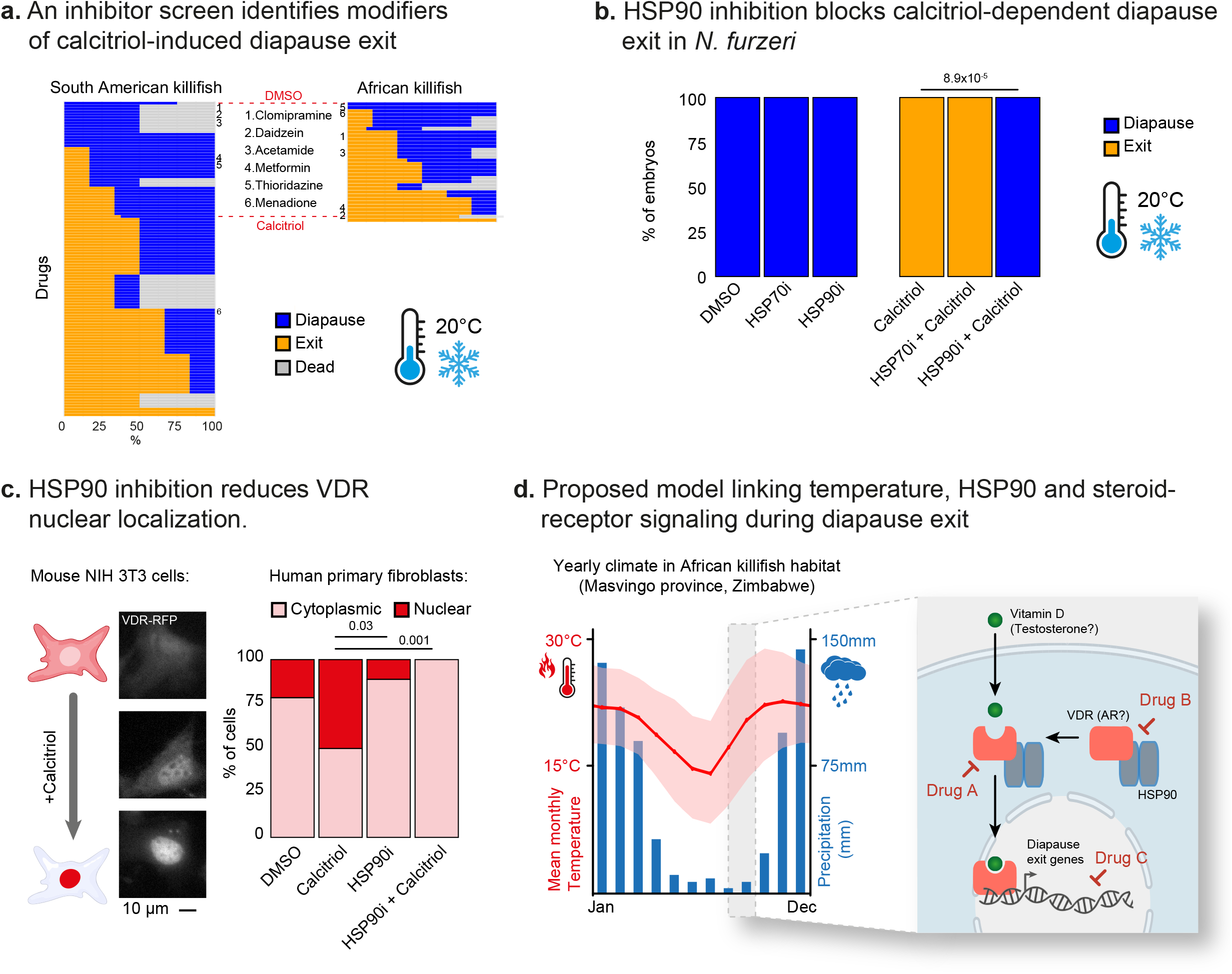
Model-based temporal transcriptomics of calcitriol-dependent diapause exit in the African killifish. **a.** Schematically illustrating the parametric functions used to model temporal transcriptome dynamics: constant (black), single sigmoid (yellow), and double sigmoid (blue). **b.** K-means clustering of temporal gene expression profiles during diapause exit. Each colored line represents the average of model-fitted values (fold change relative to time 0) for all genes in the cluster. Cluster sizes: n = 86 (1), 802 (2), 1,337 (3), 857 (4), 691 (5), 404 (6), 164 (7), 612 (8), 268 (9). **c.** Functional enrichments (rows) for each cluster. Clusters (columns) are ordered by temporal onset. Pathway enrichment was assessed at FDR < 5%. Color indicates adjusted p-value; circle size represents the number of genes associated with the pathway. **d.** Temporal expression profiles of selected genes from each cluster: *rarb* (*LOC107379177*), *cox8a* (*LOC107391695*), *eif2d* (*eif2d*) and *rarg* (*rarg*). Solid lines show measured gene expression values as log₂ fold change relative to time 0 (y-axis) over time (x-axis); shaded areas indicate the cluster average ± standard deviation. **e-f.** Transcriptome-based stochastic clock analysis estimates predicted transcriptomic age/noise structure during diapause entry (**e**) and exit (**f**). Analysis of diapause exit from this study (**f**) or entry by Hu et al.^21^ (**e**). Both predictions were generated based on training on the pre-diapause data (cream). Color gradient represents the transition from pre-diapause (cream) to diapause (blue) in (**e**), or from diapause (blue) to exit (cream) in (**f**). A sample of embryos that never entered diapause (escape) are marked in red.

Thus, by combining high-resolution single-embryo transcriptomics with parametric temporal modeling, we effectively harnessed the temporal depth of our transcriptomics data to capture the fine-grained gene expression dynamics during a vitamin D_3_-induced diapause exit trajectory and accounted for the inherent embryo-to-embryo variability of the response.

### A multi-step transcriptional cascade accompanies vitamin D₃-dependent diapause exit

We identified 5,221 genes (49%) with temporal changes in their expression during diapause exit (**Table S2**). Most of these genes (4,645, 89%) showed a single temporal change, including 4,071 up- and 574 down-regulated genes. Another 576 genes (11%) showed a transient up-(263 genes) or down-(313 genes) regulation. We used model fits to cluster all 5,221 temporally changing genes (**Figure 2b**, **Methods**). This analysis revealed 9 coherent groups with distinct temporal patterns, highlighting a multi-step transcriptional cascade of upregulated and downregulated processes. Broadly, upregulated genes could be divided into early (cluster #1), intermediate (clusters #2, 3, and 4), and late responses (clusters #5, 6, and 7). Downregulation was either transient (cluster #9) or persistent (cluster #8).

Pathway analysis (see **Methods**) provided a global overview of the cellular processes within each cluster (**Figure 2c** and **Table S3**). The early response (cluster #1) was enriched with transcription factors (TFs) and membrane transporters (**Figure 2c**). Specific genes in this cluster included steroid-hormone pathway components (e.g., androgen receptor, or ar), retinoic acid receptors (e.g., *rarb*), and their co-factors (e.g., *growth-regulating estrogen receptor binding1*, or *greb1*, **Figures 2d, S2c**). It also included immediate vitamin D_3_-responsive genes (e.g., *slc20a1*^34^), metabolic regulators (e.g., *tfeb*, a primary regulator of nematode worm diapause^35^), members of the integrated stress response (e.g., *atf4, foxo1*), growth- and proliferation-associated factors (e.g., *wnt7a, igf2, igfbp1*), and unfolded protein response genes (e.g., *crebrf*).

Intermediate upregulated genes (clusters #2, 3, and 4) were enriched for a different set of TFs, cellular respiration (e.g., *cox8a*, **Figure 2d**), and protein homeostasis (e.g., translation and degradation, *eif2d*, **Figure 2d**). Late response genes (clusters #5, 6, and 7) were associated with cell-cycle entry, reactivation of classical developmental processes (e.g., eye and somite development), and signaling pathways (e.g., *rarg*, **Figure 2d**). Finally, down-regulated clusters (clusters #8, 9) were associated with autophagy, cytokine and hormone production (e.g., insulin), developmental transcription factors (e.g., *hoxb2*), and circadian regulators (e.g., *cry1*, **Table S3**).

To explore regulatory relationships within the transcriptional program, we tested whether transcription factors upregulated in one cluster showed enrichment of their putative binding sites in the promoters of genes from a temporally subsequent cluster using HOMER^36^ (**Methods**, **Table S4**). While specific motifs were significantly enriched in our analysis (**Table S4**), none corresponded to transcription factors that were upregulated in earlier clusters. Nonetheless, focusing on smaller gene subsets within each cluster may still yield biologically meaningful insights. For example, 414 genes (46%) in cluster 2 (intermediate-response genes) contained a putative ESRRA binding site (**Figure S2d**), an estrogen signaling–associated factor. While this motif was not significantly enriched, this observation suggests the association of steroid hormones in diapause exit^37^.

Taken together, these results reveal a temporal cascade of transcriptional events underlying vitamin D₃-dependent diapause exit in African killifish, ultimately enabling embryos to terminate diapause and resume normal development.

### A stochastic clock analysis suggests reduced transcriptomic noise during diapause exit

The extended period of embryonic dormancy during diapause is not linked to significant adverse effects on adult physiology^21^. Animals that exit diapause resume normal development, exhibit typical adult lifespans, and retain full reproductive capacity. Thus, it remains unclear how embryos mitigate the cellular damage associated with prolonged dormancy, whether by limiting its accumulation during diapause, reducing it during diapause exit, or both.

To address this question, we examined transcriptional noise signatures in our data by applying a transcriptome-based clock^38^. This clock captures stochastic changes in transcriptomic noise, which have been linked to age-associated signatures; related approaches have recently been applied to embryonic dormancy in worms, human aging, and iPSC reprogramming^39^. To generate the clock, we used a previously published dataset that sampled embryos both before (pre-diapause) and during diapause^21^.

Analysis of these data using the clock suggested that transcriptomic noise increased during diapause compared to pre-diapause and escape states (**Figure 2e**), supporting the idea that noise accumulates during dormancy. Applying this stochastic clock to our diapause-exit time course revealed a decrease in predicted transcriptomic noise during exit, beginning before the major wave of cell proliferation (**Figure 2f**), suggesting that diapause exit is associated with reduced transcriptomic noise. Transcriptomic noise may decline during diapause exit as embryos become more developmentally synchronized (see **Discussion**).

### Identifying candidate pharmacological modulators of diapause exit

With vitamin D_3_ emerging as a trigger of developmental reactivation, we aimed to identify additional compounds with potential effects on diapause exit. We first compared our transcriptomic signatures (clusters) to the DsigDB^40^ drug signature database, which integrates gene-expression profiles with chemical compounds. This analysis associated 440 compounds with gene clusters of diapause exit in the African killifish (**Table S10**).

Next, we aimed to restrict this large set of compounds based on the similarities we identified in the vitamin D_3_ trigger between exit and escape from diapause, and between the African and South American killifish clades. We therefore applied a similar analysis to published transcriptomic data from *A. limnaeus* embryos^17^ (**Figure 3a**). These data were collected at comparable developmental stages (preceding diapause) in two trajectories: temperature-dependent escape from diapause (30°C) or entry into diapause (20°C). PCA analysis of these data confirmed that PC1 captured developmental progression in the expected developmental order (**Figure 3b**, left), while PC2 separated the two developmental trajectories. Interestingly, although overall developmental stages are comparable, commitment to each trajectory is clearly separated by PC2 in even the earliest of samples, which supports previous data that developmental trajectory is determined many days prior to entry into diapause (**Figure 3b**, left). Finally, applying stochastic clock analysis (see **Methods**) to this dataset indicated a progressive increase in transcriptomic noise during both developmental trajectories (**Figure 3b**, right). RAPToR^41^-based transcriptomic staging showed a similar late-stage deviation from the escape-reference trajectory (**Figure S3a**).

**Figure 3.**
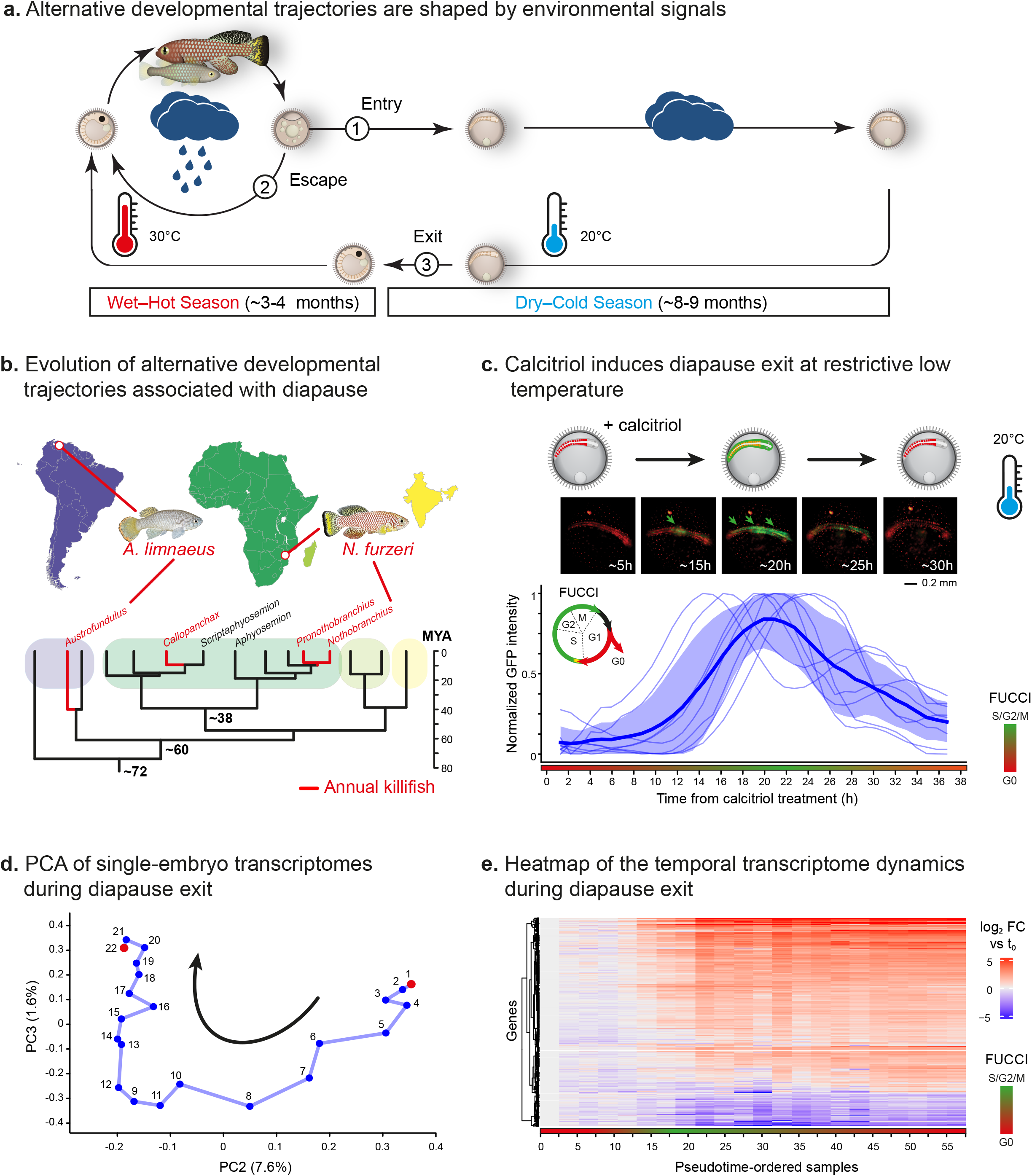
Temperature-dependent transcriptional programs associated with diapause entry and escape in *A. limnaeus*. **a.** Experimental design (Data from Romney et al., 2018^17^). Embryos were incubated at either 20 °C (to induce diapause entry) or 30 °C (to escape diapause). RNA was extracted from embryos at seven morphological stages: DC (dispersed cell), NK (neural keel), 6S (6 somites), 10S (10 somites), 16S (16 somites), 20S (20 somites), and 24S (24 somites). Data from Romney et al., 2018^17^. **b.** Left: PCA of RNA-seq (Data from Romney et al., 2018^17^). Each dot represents one replicate collected at the indicated embryonic stages, for either escape (shades of red) or diapause (shades of blue). Right: Transcriptome-based stochastic clock analysis estimates transcriptomic noise during escape (red) or diapause (blue; see **Methods**) trajectories. **c.** Gene clusters grouped by differential expression patterns between the two conditions: 30 °C (escape, red) and 20 °C (diapause, blue). Clusters are divided into two rows based on gene expression: higher expression in escape (top) or in diapause-promoting conditions (bottom). Clusters are divided into 6 columns by dynamic behaviors: increase–increase, increase–constant, increase–decrease, constant–constant, constant–decrease, or decrease–decrease. Each panel shows the average gene expression in the cluster. All data were normalized to the mean expression of both conditions at the first developmental stage. The number of genes in each cluster is indicated in brackets.

We adapted our temporal modeling framework and applied it to the *A. limnaeus* data to compare gene expression between the two conditions and identify genes with trajectory-specific expression differences, arising either from distinct dynamics or levels (**Methods**). This analysis revealed 2,954 genes (42%) with trajectory-specific dynamics or level differences between the diapause-entry and escape trajectories (**Tables S5-7**). We classified these genes into 12 groups (groups #1.1–6.2, **Figure 3c**), defined by their dynamics (upregulated, constant, or downregulated; columns in **Figure 3c**) and relative expression levels (higher in one trajectory versus the other; rows in **Figure 3c**). The remaining 58% of genes (4,046 genes, **Figure S3b, Table S8**) displayed similar expression in both trajectories and were therefore interpreted as representing other transcriptional programs not directly involved in diapause regulation (**Figure S3c**).

Functionally, genes with higher expression in the escape trajectory were associated with energy production and cellular respiration (e.g., *cycs*), protein folding (e.g., *hsp90b1*), vitamin D_3_ signaling, and mRNA processing and translation (**Figure S3d**, **Table S9**). In contrast, genes with higher expression in the diapause trajectory were associated with organelle disassembly and autophagy (e.g., *becn1*). A comparison to the African killifish transcriptomics recovered shared transcriptional modules despite substantial differences in trigger, developmental context, and timescale (**Figure S3f**).

We then applied the same DSigDB drug-enrichment analysis to the trajectory-specific gene clusters identified in the South American killifish dataset. This analysis associated 496 compounds with diapause entry/escape signatures (**Table S10**). Integrating the African and South American killifish drug-enrichment outputs yielded an approximately 600-compound transcriptome-guided candidate set, including 291 compounds predicted in both species (**Figure 4a**, left). We prioritized this cross-species overlap because these compounds are more likely to target shared diapause-regulatory modules rather than species-or stimulus-specific responses.

**Figure 4.**
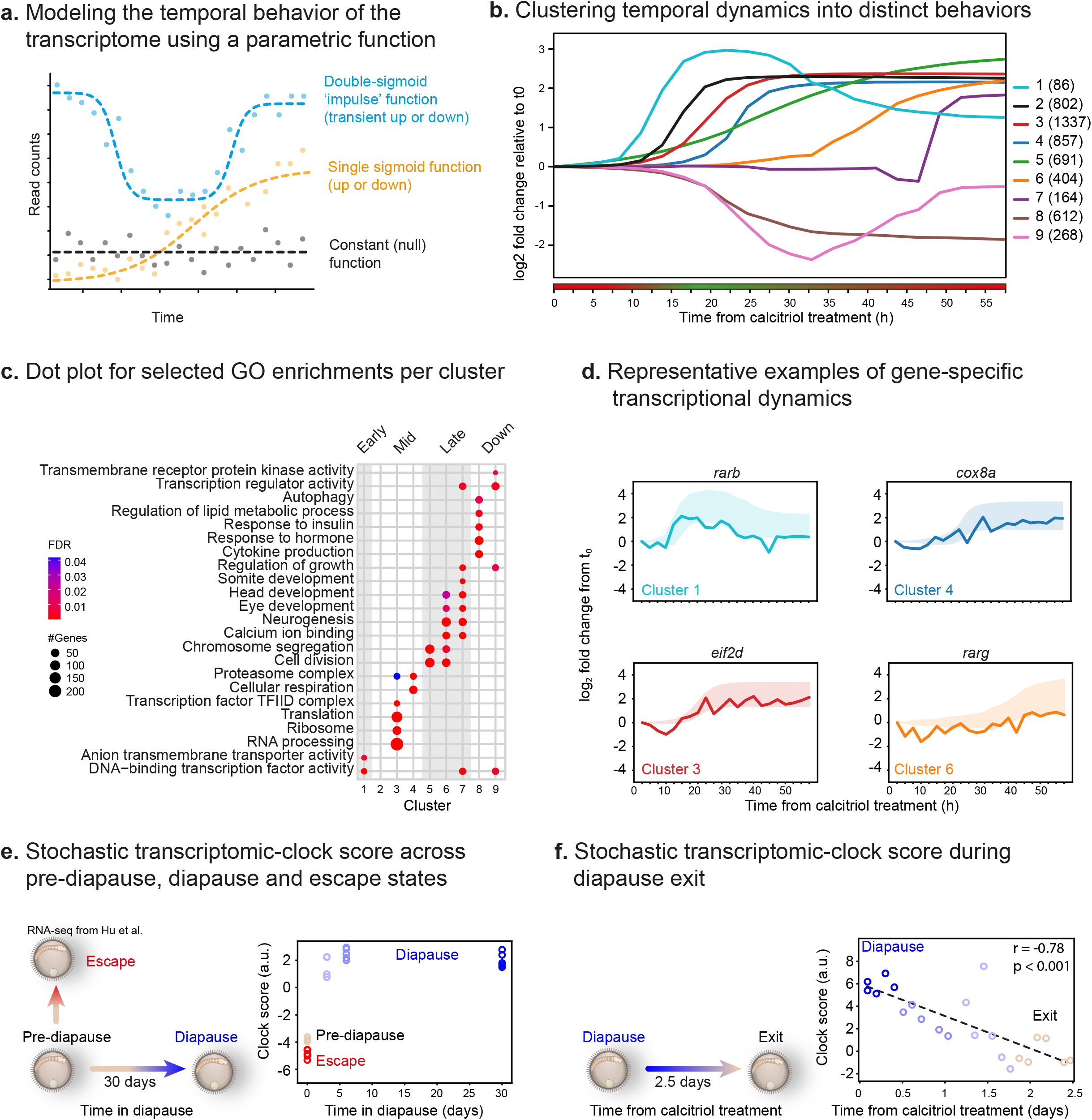
Transcriptome-guided small-molecule screening reveals shared regulators of diapause exit across killifish clades. **a.** Drug-enrichment analysis was performed on transcriptomic clusters from each species using DSigDB. Compounds predicted in both species were prioritized for functional testing as candidate conserved modulators of diapause-related transitions (**Left**). For each subgroup, one representative compound is highlighted in red, and its molecular structure is displayed as an example of the group’s chemical characteristics (**Right**). **b.** Experimental design for the chemical screen to identify drugs that promote diapause exit (activator screen) or inhibit the calcitriol-induced exit from diapause (inhibitor screen). The complete 111-compound library was screened in *A. limnaeus*, and selected compounds were subsequently tested in *N. furzeri*. **c.** Exit from diapause can be triggered by exposure to 1 μM testosterone (African killifish: DMSO (2%), n=37; calcitriol (100nM), n=41; testosterone, n=19; South American killifish: DMSO (2%), n=20; calcitriol (100nM), n=23; testosterone, n=24) in both killifish clades. Bar graphs represent the proportion of embryos (y-axis) that either exit (orange) or maintain diapause (blue) following treatment as indicated (x-axis) at 20 °C. P-values are shown above, calculated by a Fisher’s exact test with 5% FDR correction.

Compounds identified in both species were linked to various physiological processes (**Figures 4a**, right, **S4a**), such as steroid hormones (e.g., testosterone), neuronal and immune system modulators (e.g., clomipramine and cyclosporin A, respectively), and metabolic regulators (e.g., metformin).

Together, these results identify candidate chemical modulators of diapause-related state transitions that are likely independent of clade and upstream cues (calcitriol or temperature).

### A targeted chemical screen identifies activators and inhibitors of diapause exit

Diapause regulation may repurpose a core developmental network. Consequently, functional analysis requires reversible manipulation within a defined developmental window, limiting the usefulness of conventional irreversible genetic perturbations (e.g., CRISPR). Moreover, candidate-based genetic approaches are inherently low-throughput, and previous attempts have achieved only partial reversal of the diapause phenotype^21,42^. We therefore used the transcriptome-guided pharmacological predictions described above to assemble a single targeted small-molecule library for functional testing. Similar approaches have been instrumental in revealing developmental mechanisms in zebrafish^43–48^.

From approximately 600 predicted compounds, we selected a final set of 111 compounds for a single targeted screen (**Tables S11-13**). Sixty-nine compounds were selected directly from the drug-enrichment analysis, prioritizing compounds with stronger enrichment scores, cross-species support, and availability in aliquoted screening-ready format. An additional 42 compounds were included based on biological relevance to enriched pathways and candidate regulatory nodes, including steroid/nuclear-receptor signaling, metabolism, AKT/mTOR/FOXO signaling, and protein-homeostasis. We excluded a subset of proliferation-targeting chemotherapeutic, highly toxic, or poorly stable compounds.

To test these compounds as activators or inhibitors of diapause exit (see **Methods**), we used a simplified strategy that decoupled temperature effects from vitamin D_3_ regulation. Diapausing embryos from both clades were incubated in 96-well plates at diapause-inducing temperature (20°C) with 1 µM of each selected compound (6–9 embryos per compound) for 3 weeks, and monitored for eye emergence as an indicator of diapause exit (**Figure 4b**, **Tables S11-13**). The complete 111-compound library was screened in *A. limnaeus*, and a selected subset was subsequently tested in *N. furzeri*.

We designed two complementary screening strategies. In the activator screen, compounds that mimicked calcitriol by inducing diapause exit at 20°C were considered hits (≥25% of treated embryos exited diapause; see **Methods**). In the inhibitor screen, compounds that reduced calcitriol-induced diapause exit by ≥50% relative to the calcitriol-only control were classified as potential inhibitors We screened the final set of 111 compounds using both the activator and inhibitor strategies. The activator screen revealed that testosterone, the primary steroid sex hormone in males, and its synthetic agonist (i.e., danazol^49^), are effective in inducing diapause exit in both killifish clades (**Figure 4c**, independent replication in **Table S14**, and see previous studies involving female sex hormones^37^). Notably, testosterone and danazol emerged as the primary robust activators from this targeted library.

In contrast, the inhibitor screen identified a more diverse set of compounds that blocked calcitriol-induced diapause exit with varying efficiencies in the South American killifish (**Figure 5a**, **Tables S12-13**). These included proliferation inhibitors (e.g., gemcitabine, doxorubicin, and etoposide^50^), psychiatric drugs (clomipramine), nonsteroidal anti-inflammatory drugs (e.g., piroxicam), and nuclear receptor agonists (e.g., pirinixic acid^51^). Application of the inhibitor screen in the African killifish with a subset of 30 selected compounds (**Table S13**) confirmed a functional conservation for 46.6% (14/30) of tested compounds (**Figure 5a**), including metabolic regulators^52^ (metformin) and antidepressants (clomipramine).

**Figure 5.**
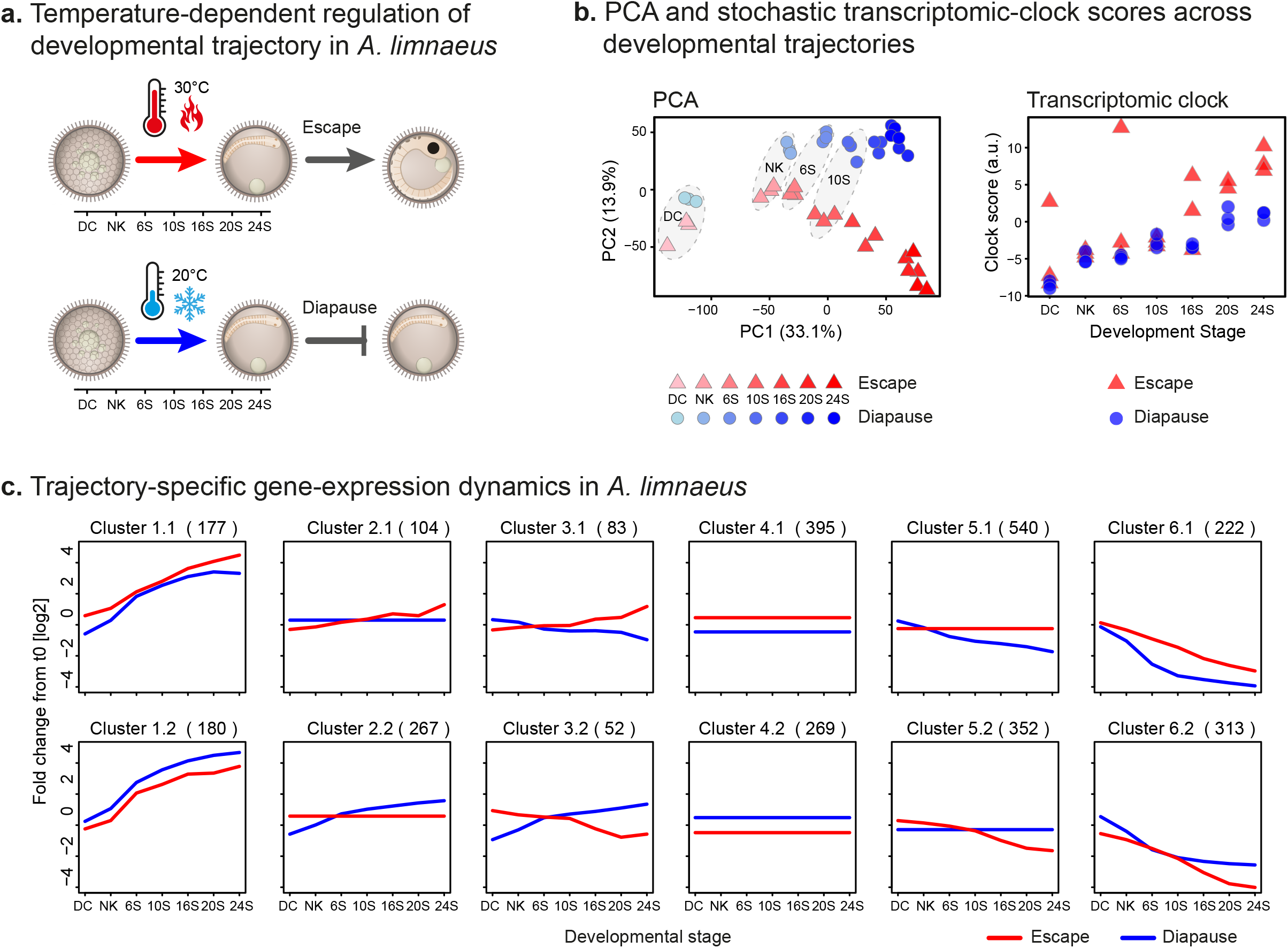
HSP90 integrates temperature cues with vitamin D₃ signaling to regulate diapause exit. **a.** Inhibitor screen identifies modulators of calcitriol-dependent diapause exit. Results of drug screen in the South American killifish (left) and follow-up testing of a selected subset in the African killifish (right). The proportion of embryos (x-axis) in diapause (blue), exit state (orange), or dead (grey) is indicated for each of the tested compounds (y-axis). Positions on both graphs are shown for six selected drugs (numbered 1-6). As a positive control, calcitriol was added alone, and DMSO (without calcitriol), was used as a negative control (see red dashed lines). Dead embryos are shown for completeness but were excluded from calculation of exit rates and statistical analyses. **b.** Pharmacological inhibition of HSP90 blocks calcitriol-dependent diapause exit in African killifish. Bar graphs represent the proportion of embryos (y-axis) that either exit (orange) or maintain diapause (blue) following treatment as indicated (x-axis) at 20 °C after one week. P-values from Fisher’s exact tests are shown. DMSO (3%); HSP70i (200µM); HSP90i (200µM); calcitriol (100nM); calcitriol (100nM) + HSP70i (200µM); calcitriol (100nM) + HSP90i (200µM). n=8-12 (see **Methods**). **c.** Left: representative images from transfection of a VDR–RFP construct into murine NIH3T3 cells, Scale bar 10µm. Right: transfection of the same construct into primary human fibroblasts and quantification of VDR nuclear translocation from the cytosol (pink) to the nucleus (red). Bar graphs represent the proportion of cells (y-axis) with either a cytoplasmic (pink) or nuclear (red) localization of VDR-RFP following treatment as indicated (x-axis). P-values are calculated by Fisher’s exact test with 5% FDR correction. For NIH3T3 source data and statistics, see **Source Data** and **Methods**. For Human fibroblast, 14-30 cells were imaged per condition, and. Experiments were replicated twice. **d.** HSP90 links environmental seasonal temperature rise to steroid-dependent diapause exit. Left: Climatological data showing yearly temperature and precipitation in Masvingo Province, Zimbabwe, a primary natural habitat of *N. furzeri* (adapted from Climate-Data.org). Right: Schematic model of the thermo-sensory signaling axis. Elevated environmental temperatures support a model in which HSP90’s enhanced chaperone activity is required for the proper folding and ligand-binding capacity of steroid hormone receptors. Small-molecule inhibitors that successfully blocked diapause exit are mapped onto the vitamin D₃ pathway, acting as predicted modulators of the steroid receptor (Drug A), the HSP90 chaperone complex (Drug B), or downstream effectors (Drug C). Climate data based on Climate-Data.org.

Mapping compounds that blocked diapause exit onto the vitamin D_3_ signaling pathway revealed multiple potential targets, particularly drugs affecting the interaction between vitamin D_3_ signaling and the AKT/mTOR/FOXO axis^53,54^ (e.g., the AKT inhibitor afuresertib^55^, the FOXO inhibitor diethyl maleate^56^, and the antidiabetic and mTOR-interacting metformin^57–59^). Other small molecules were modulators of steroid receptors (e.g., clomipramine^60^ and daidzein^61^) and of the heat shock protein HSP90 (e.g., acetamide^62^, clomipramine^63,64^, daidzein^65^, and metformin^66^).

These results establish a tractable small-molecule platform for perturbing diapause exit and identify both shared and species-specific pharmacological regulators. Rather than nominating individual compounds as definitive pathway-specific probes, the screen highlights central regulatory nodes that can be tested mechanistically.

### HSP90 links temperature and vitamin D_3_ signaling during diapause exit

The temperature-dependent activation of protein homeostasis (proteostasis) during the escape trajectory (e.g., ‘protein folding’, **Figure S3d**) suggested that the heat-shock response might act as an integrator of environmental cues, linking vitamin D₃ and temperature (e.g., ‘VDR pathway’, **Figure S3d**). Consistently, several HSP90-related genes^67^ were upregulated during the temperature-induced escape trajectory in the South American killifish (e.g., *stip1*, *ptges3*, *ppid*, **Figure S5a**, and *hsp90b1*, **Figure S3e**). Because morpholino knockdown is unlikely to remain stable through the prolonged killifish diapause-synchronization period, we used acute, reversible pharmacological perturbation to target the specific window of diapause exit.

We exposed diapausing embryos from both clades to inhibitors of HSP90 (HSP90i, ganetespib^68^) or HSP70 (VER-155008^69^) and tested whether these perturbations block calcitriol-induced exit (**Figures 5b, S5b**; **Table S14**). HSP90 inhibition prevented calcitriol-induced exit after one week at 20°C, whereas HSP70 inhibition did not. HSP90 inhibition showed a trend toward reduced diapause exit at higher temperature in the African killifish (although this effect did not reach statistical significance, p = 0.06, 30°C), where lower doses of calcitriol are sufficient to induce exit (**Figure S5c**; **Table S14**). The inhibitory effect weakened after two weeks, suggesting either delayed calcitriol action or limited ganetespib stability. These data identify HSP90 activity as a functional requirement for efficient calcitriol-dependent exit from diapause.

Previous studies in cancer cells^70,71^, primarily using shRNA, demonstrated that, similar to other steroid receptors (e.g., the androgen receptor), maturation and activation of the vitamin D receptor (VDR) depend on proper folding mediated by HSP90. We therefore hypothesized that HSP90 inhibition impairs diapause exit by limiting VDR maturation and nuclear signaling, and tested whether ganetespib reduces calcitriol-induced VDR nuclear translocation in non-cancerous cells. Indeed, in primary human fibroblasts grown in steroid-depleted serum and transfected with RFP-VDR^72^, acute calcitriol stimulation triggered nuclear accumulation that was significantly reduced by HSP90 inhibition (**Figure 5c**; **Table S15**). We observed the same effect in murine NIH3T3 cells (p = 0.022; **Table S16**).

Together, these data support HSP90-dependent VDR trafficking as a plausible mechanism linking steroid signaling to diapause exit. Consistent with a shared role across clades, HSP90 inhibition impaired calcitriol-induced exit in both killifish species. This proposed mechanism allows the integration of environmental cues, particularly the rise in temperature that precedes the rainy season (**Figure 5d**), into the molecular networks that regulate embryonic dormancy.

## Discussion

In this study, we show how environmental and hormonal cues are integrated into the molecular circuits that control exit from diapause and the resumption of development. By combining high-resolution single-embryo transcriptomics with chemical perturbation screens, we overcome the inherent variability of these programs and define a regulatory framework in which HSP90 provides a mechanistic link between environmental conditions and nuclear-receptor signaling during developmental reactivation in killifishes. A cross-clade analysis of the molecular network and pharmacological responses extends the generality of this framework.

### A shared regulatory framework for dormancy exit

The results and analyses of this study suggest that core modules in the diapause program are shared across related annual killifish clades. These clades shared drug-enrichment signatures in transcriptomic data, vitamin D_3_-induced diapause exit, pharmacological modulators identified in the chemical screen, and HSP90 modulation of diapause exit. A comparative analysis of the transcriptomic datasets used in this study further suggests that core modules might also be shared across clades. These may represent reusable regulatory elements that remain active across multiple diapause-related transitions, including modules for cell-cycle control, autophagy/proteostasis, and metabolic remodeling. Still, our cross-clade comparisons remain limited by differences in trigger and developmental context: diapause exit versus entry and escape, calcitriol-versus temperature-induced, and different staging schemes and timescales. Future studies directly measuring diapause exit across clades will be needed to confirm these observations.

The evolutionary origin of this shared regulatory machinery remains unresolved. Diapause may have evolved independently in African and South American annual killifishes, as previously proposed^22,23^, or may have been present in a dormancy-capable common ancestor and subsequently retained or elaborated in annual lineages; the current study cannot distinguish between these alternatives.

The latter possibility is consistent with observations that semi-annual killifishes can exhibit diapause-like states under certain conditions. Future experiments applying our approach to non-or semi-annual outgroups could distinguish between these evolutionary scenarios.

Shared regulators such as TFEB (**Figure S2c**), previously implicated in dauer regulation in *C. elegans* (possibly via *TGF-β*^35,73^), further support continuity between dormancy-associated control programs across phyla. The repeated engagement of proteostasis-related pathways in both killifish clades also suggests that diapause transitions draw on stress-response modules with broader relevance to development and aging^74^. It is worth noting that differences in exit timing that appear stochastic could result from embryo-to-embryo molecular differences in baseline factors that drive calcitriol-response thresholds.

### A platform for small-molecule screening in killifish diapause

Measuring and modeling interventions to manipulate physiology is challenging^40,75^. The screen was therefore not an unbiased whole-library screen, but a transcriptome-guided perturbation assay designed to test whether diapause-associated gene-expression modules nominate conserved pharmacological nodes. The overlapping drug–gene signatures across clades reinforce the usefulness of pharmacological perturbation as a discovery tool. We interpret the screen mainly as a way to identify central regulatory nodes rather than to validate individual compounds. Under that framework, the convergence of chemically diverse perturbations on HSP90 is especially informative. Thus, integration of vitamin D_3_ and temperature through common downstream modulators such as HSP90 provides a plausible mechanistic solution for coupling environmental state to developmental phenotype. The additional positive hits from the screen likely point to a wider network of modulators that remains to be resolved.

### Coupling environmental sensing to developmental timing

Successful diapause exit must be timed with exceptional precision: if embryos reactivate too early or too late, offspring may encounter conditions they cannot survive. Our findings, therefore, point to a broader developmental principle, in which molecular circuits control developmental timing by translating environmental information into coordinated cell-state transitions. In this context, the HSP90-linked network implicated here may represent one component of a broader adaptation that couples environmental conditions with developmental decisions, allowing vertebrates to couple environmental sensing with developmental decisions across life-history transitions. The conceptual and experimental methodology developed in this study may motivate further studies on resilience mechanisms under climate change. Crucially, as environmental instability and warming intensify, these findings suggest how climate change may impact embryogenesis. By identifying a molecular mechanism contributing to environmental sensing and developmental reactivation, this framework may help guide future studies of how changing environmental conditions influence embryonic developmental decisions and resilience.

### HSP90 is candidate integrator of environmental cues for diapause exit

We identify the HSP90 chaperone as an integrator in the temperature-sensing network, well positioned to translate environmental inputs into developmental outcomes. Indeed, across eukaryotes, including yeast, plants^76^, and animals such as Mexican cavefish^77^ and humans^78^, HSP90 supports growth and development by stabilizing a wide range of client proteins. These clients include components of insulin/IGF signaling^79,80^, a pathway already implicated in diapause regulation in both worms^81^ and killifish^15^ (as well as in aging^82^). By promoting the proper folding and stability of client proteins, HSP90 buffers against genetic and environmental variation, a stabilizing role observed in a variety of species including zebrafish^83,84^. However, this buffering capacity can be overwhelmed by extreme environmental stress, such as that endured by annual killifishes. When this capacity is breached, hidden genetic variation is unmasked, allowing marked phenotypic shifts to emerge even in the absence of strong external cues^83^. Consistent with this idea, several HSP90-related genes exhibit contrasting evolutionary signatures between annual and non-annual killifishes. For example, *hsp90b1,* and the HSP90 cofactors *CDC37* and *FKBP4*, show intensified selection in annual killifishes but relaxed selection in non-annual killifishes^85^. Together, these patterns suggest that distinct selective pressures in annual versus non-annual killifish lineages have differentially shaped vitamin D_3_ and HSP90-associated pathways.

### Transcriptomic noise during diapause exit

Killifish diapause represents an extreme form of lifespan plasticity: embryos can remain developmentally arrested far longer than the usual lifespan of the species, yet later resume development with apparently normal growth and reproduction. Our transcriptome-based clock^38^ analysis suggests that age-associated noise is accumulated during diapause, but this trend is reversed once embryos exit diapause and resume development (**Figure 2f**). Decreasing transcriptomic noise during diapause exit may reflect increased synchrony among embryos as part of developmental re-entry or a return toward a developmental ground state. Related evidence from previous work^86^ shows distinct ubiquitin-conjugate dynamics in diapause II and post-diapause embryos, consistent with remodeling of proteostasis across this transition. The accumulation of ubiquitin conjugates during anoxia is consistent with the possibility that proteasome activity is more strongly suppressed in diapause II embryos than in post-diapause embryos, suggesting increased protein turnover following developmental reactivation. These observations motivate the hypothesis that diapause exit may involve reversal of aging-relevant molecular damage.

It is worth noting that while our temporal clustering defines the ordered gene-expression program of diapause exit, the transcriptomic aging clock estimates age-associated stochastic transcriptional noise rather than organismal age itself. Moreover, the clock data for the South American killifish suggest that transcriptional noise is more strongly aligned with developmental stage than with temperature (particularly, up until the 10-somite stage, **Figure 3b**), suggesting that temperature has a relatively modest effect in this dataset. Nonetheless, we cannot rule out that temperature may confound the South American killifish comparison, as faster clock progression could reflect temperature rather than developmental trajectory. Finally, although the analysis met the computational pipeline’s validity criteria, the limited clock-training set led us to present these findings as exploratory rather than definitive evidence of age reversal.

As an orthogonal developmental-staging analysis, RAPToR showed a qualitatively similar late-stage transcriptomic realignment (**Figure S3a**). This analysis revealed a decline in developmental stage at advanced pre-diapause stages, most prominently around 20–24 somite stages, closely matching the stochastic BiT-Age clock results (**Figure 3b**), and providing independent support that both approaches capture the same diapause-associated transcriptomic realignment. As diapause exit in the African killifish is largely morphologically static over the 72-hour window, this analysis could be applied only to the South American killifish dataset, using escape-trajectory embryos as the developmental reference to predict developmental stage in the diapause trajectory.

Interpreting the observed transcriptomic noise signatures is more challenging. Several explanations remain possible. Noise could be reduced during diapause exit due to (1) an age-associated reversal driven by restorative processes during diapause exit; or (2) rapid transcriptional resynchronization associated with the resumption of development. On the other hand, increased transcriptional noise during diapause may arise from (1) stochastic transcriptional changes during diapause caused by incomplete synchrony among embryos, consistent with diapause not being a fully static state; or (2) mechanisms that globally repress transcription during diapause.

One intriguing possibility is that reduced transcriptional noise reflects age-associated reversal driven by restorative processes during diapause exit. While earlier work in *C. elegans*^39^ and studies benchmarking stochastic transcriptomic clocks in aging^39^ and reprogramming^87^ provide useful context, these results should still be interpreted with caution due to the specialized state that diapause represents. Because this metric quantifies age-associated transcriptomic noise rather than organismal age itself, these results most likely provide evidence for reversal-like transcriptional remodeling and a possible developmental restart, rather than definitive proof of rejuvenation. Orthogonal approaches, such as quantification of damage reversal during exit, could provide independent support for this hypothesis. Loss of proteostasis is a central hallmark of aging and ubiquitin-dependent protein turnover declines or becomes dysregulated with age^88^. Our previous work^86^ showed that diapause II and post-diapause embryos differ markedly in ubiquitin-conjugate dynamics during stress and recovery: diapause II embryos accumulate ubiquitin conjugates during stress, whereas 4-day post-diapause embryos do not, but instead show increased ubiquitin conjugates during recovery. These data suggest that diapause and diapause exit are associated with distinct proteostasis states. This raises the testable hypothesis that diapause exit involves coordinated remodeling of aging-relevant stress-response pathways, including proteostasis. Future studies using orthogonal markers, such as protein aggregation, DNA damage, oxidative damage, mitochondrial function, senescence-like signatures, and epigenetic marks, will be required to determine whether diapause exit involves true biological rejuvenation.

More broadly, diapause exit offers a tractable vertebrate system for dissecting how tissues re-enter development after prolonged dormancy without obvious long-term loss of function.

---

**Figure S1.**
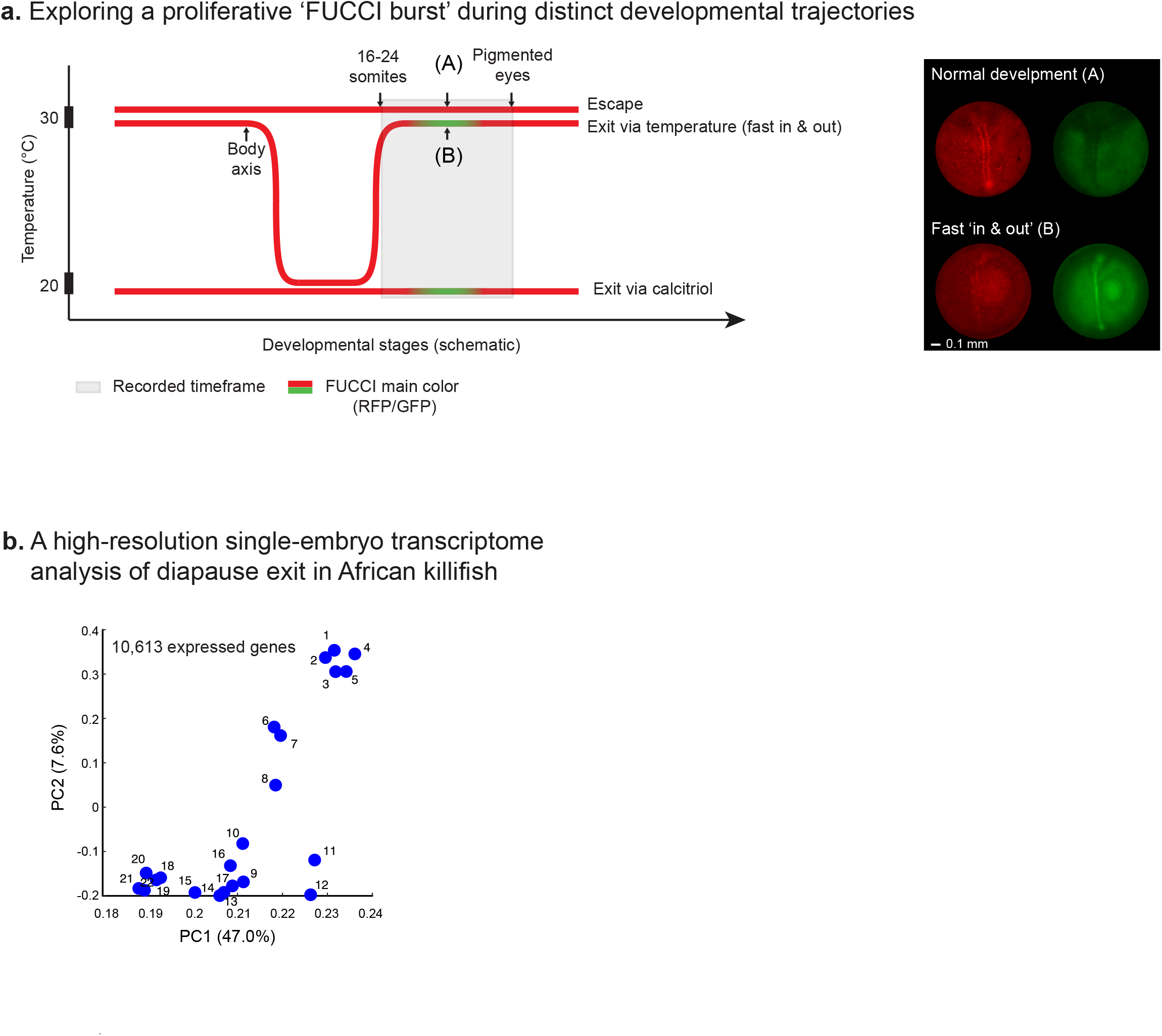
Comparative dynamics of escape and diapause exit. **a.** Left: Schematic representation of FUCCI embryo development across three trajectories: Direct development without diapause (escape, n=3); Temperature-induced exit, defined as rapid exit from a brief (2 days) diapause followed by exit triggered by thermal shift (n=10); and Vitamin D_3_-induced exit (n=55, an independent experiment from Figure 1C). Grey shaded regions indicate the recorded observation periods, and arrows mark key developmental milestones. Right: Corresponding microscopy images: (Top) An embryo raised at 30°C (escape), imaged within a developmental window in which the GFP burst is expected (between the 16-somite stage and emergence of eye pigmentation). (Bottom) RFP and GFP channels of an embryo after transfer from 20°C to 30°C (rapid exit). Scale bar: 0.1mm. **b.** PCA of single-embryo RNA-seq data (x-axis: PC1, y-axis: PC2). Each symbol represents one embryo collected at 2.5-hour intervals after calcitriol treatment, over a total duration of ∼60 hours. 10,613 genes used for PCA.

**Figure S2.**
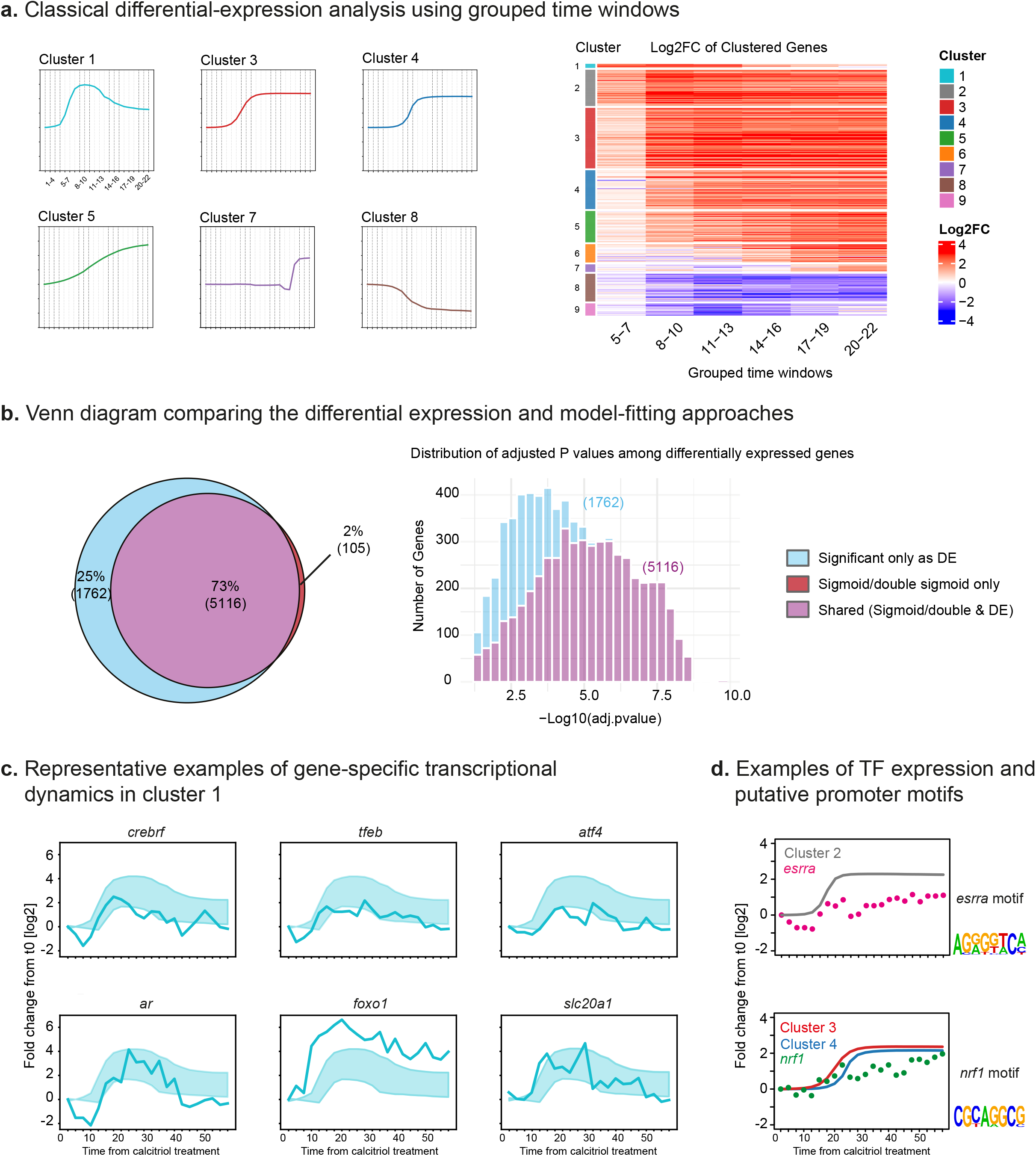
Validation of diapause exit model-based clustering and calcitriol-dependent transcriptional regulatory networks. **a)** Comparison of temporal model-based clustering and classical pairwise differential expression (DE). **Left**: Gene expression clusters corresponding to Figure 2b, annotated with seven defined time-windows (vertical stripes). The single-embryo temporal samples within each time window, representing 3-4 consecutive samples, were grouped for the purpose of performing a pairwise analysis. Time-windows consist of 3 time points each, except for the first window (4 time points). **Right:** Heatmap displaying the log2(Fold Change) of gene expression in six windows spanning timepoints 5–22 of the time course relative to the initial diapause exit baseline (timepoints 1–4). Timepoints 8–10 correspond to the FUCCI GFP burst (see Figure 1c). Genes (rows) are ordered and color-coded according to the model-based clustering. **b)** Overlap between model-based and pairwise differential expression. **Left**: Venn diagram illustrating the intersection of genes identified by two analytical approaches: genes rejecting a constant model in favor of a dynamic differential model by temporal modeling (sigmoid or double sigmoid, Red + Purple); and genes identified as differentially expressed in at least one of 6 time-windows using a pairwise comparison against the initial window (|log2FC| > 1, adj. *P* < 0.05; Light-blue + Purple). The intersection (Purple, *n* = 5,123) represents high-confidence DE genes supported by both methods. Percentages indicate the proportion of genes identified by at least one of the two approaches. Only 2% is identified only by temporal modeling without support from classical DE analysis, while 25% were identified only by DE without support from temporal modeling, validating the high stringency of the temporal modeling approach. **Right**: Histogram showing the p-value distribution of significant genes in the standard differential expression (DE, for the light-blue and purple groups). This distribution indicates that genes captured by the sigmoid and double-sigmoid models represent a subset of the most highly significant genes identified by the DE analysis. **c.** Temporal expression profiles of selected genes from cluster 1: *crebrf* (LOC107382399), *tfeb*, a*tf4* (LOC107376364), *ar*, *foxo1(*LOC107386483), and *slc20a1*(LOC107393652). Solid lines show measured gene expression (y-axis) over time (x-axis); shaded areas indicate the cluster average ± standard deviation. **d.** Promoter motif enrichment (+100 bp to −1 kb) links transcription factors to cluster regulation. ESRRA motif is identified in 414 genes in cluster 2 (gray), with *esrra* expression (red) preceding target activation. NRF1 motif is identified in 138 genes in clusters 3 (red) and 4 (blue), with *nrf1* expression (green) showing similar timing.

**Figure S3.**
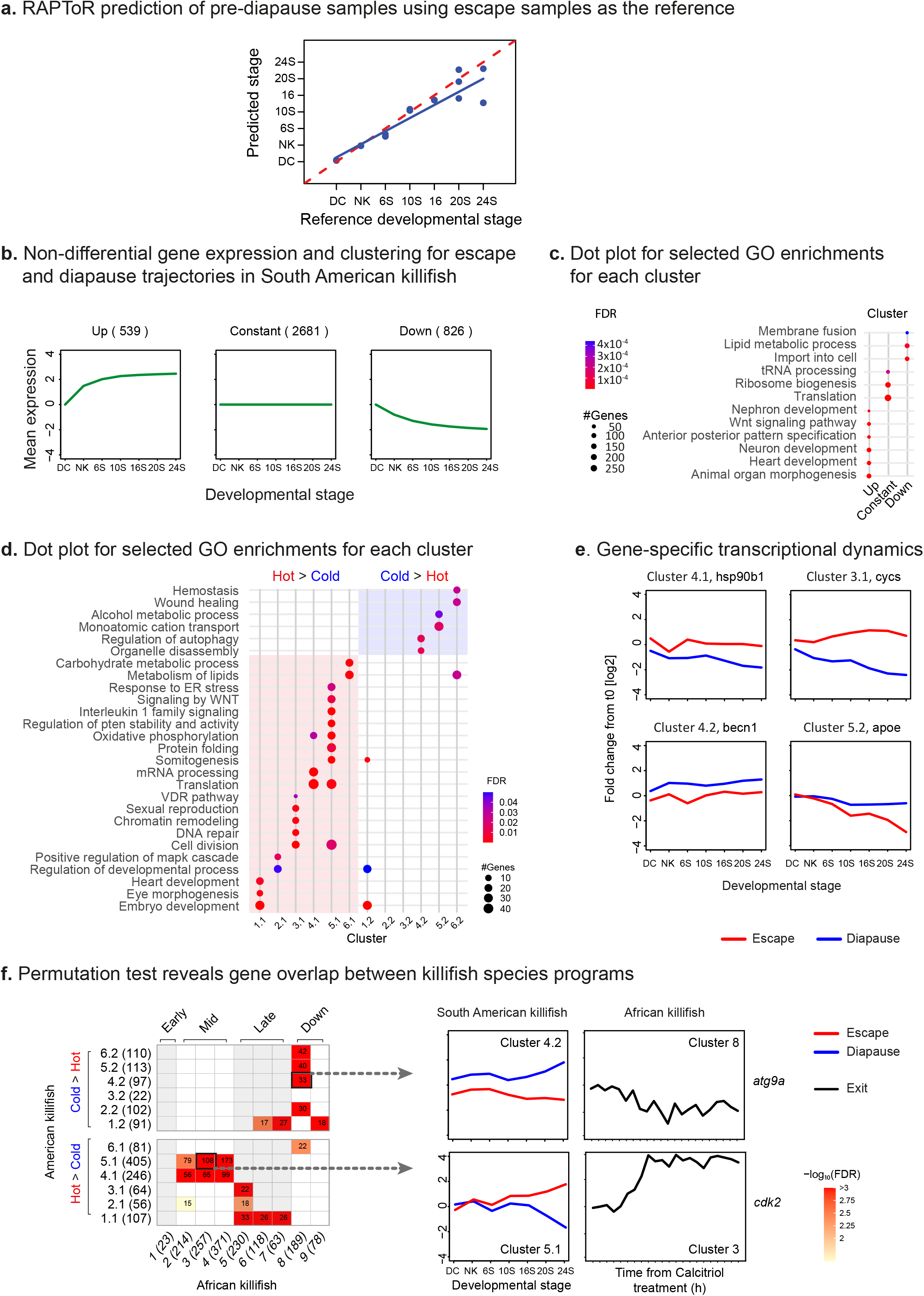
Temporal RNA-seq analysis of diapause and escape trajectories in the South American killifish. **a.** Reference developmental stage versus RAPToR^41^-predicted stage estimates in *A. limnaeus* embryos. RAPToR, a transcriptome-based developmental staging framework, requires a normal-development reference; therefore, escape-trajectory embryos developed at 30°C were used as the reference baseline across seven morphological stages: DC, dispersed cell; NK, neural keel; and 6S, 10S, 16S, 20S and 24S, corresponding to the indicated somite stages. Predicted developmental stage was then estimated for diapause-trajectory embryos developed at 20°C, shown as blue dots. The dashed red line indicates perfect agreement with the escape-reference trajectory, and the solid blue line shows the linear regression fit for the diapause-trajectory samples. **b.** Groups of non-differentially expressed genes (n = 4,046) between diapause and direct development (escape) conditions: increase (n = 539, left), constant (n = 2,681, middle), and decrease (n = 826, right). Each panel shows the mean normalized gene expression in the cluster relative to t₀ (green lines). The number of genes in each cluster is indicated in brackets. **c.** Functional enrichment (GO, KEGG, Reactome, rows) per cluster (columns) at FDR < 5%; color denotes adjusted p-value, and circle size indicates gene count per pathway. This analysis suggests that constant genes are functionally associated with housekeeping processes, such as ribosome biogenesis and translation; up-regulated genes related to developmental processes (possibly preceding the decision on diapause entry or escape), and down-regulated genes associated with lipid metabolism and cellular transport. **d.** Functional enrichments for each cluster (shown in Figure 3c). Pathway enrichment was assessed at FDR < 5%. Color denotes adjusted p-value; circle size indicates the number of genes in the pathway. **e.** Temporal expression profiles of representative genes from selected clusters. Expression values (y-axis) over time (x-axis) are shown for both conditions, normalized to the mean expression of both trajectories at the first stage (“DC”). **f.** Left: Gene overlap was assessed using a permutation-based test (Cluster sizes shown here refer to genes retained after restriction to the cross-species shared-gene set; see **Methods**). Cell values represent the number of shared genes, and color intensity reflects FDR-corrected p-value (−log₁₀ scale). Rows and columns are ordered by temporal expression patterns: *N. furzeri* clusters are ordered according to their progressive expression patterns, with downregulated clusters placed at the end. *A. limnaeus* clusters are ordered based on their expression levels in the 30°C and 20°C conditions, with clusters showing higher expression in 30°C placed lower and those with higher expression at 20°C placed higher. Right: Example genes from overlapping clusters. Gene expression is shown for diapause (blue) and escape (red) trajectories in the South American killifish, and for diapause exit (black) in African killifish.

**Figure S4.**
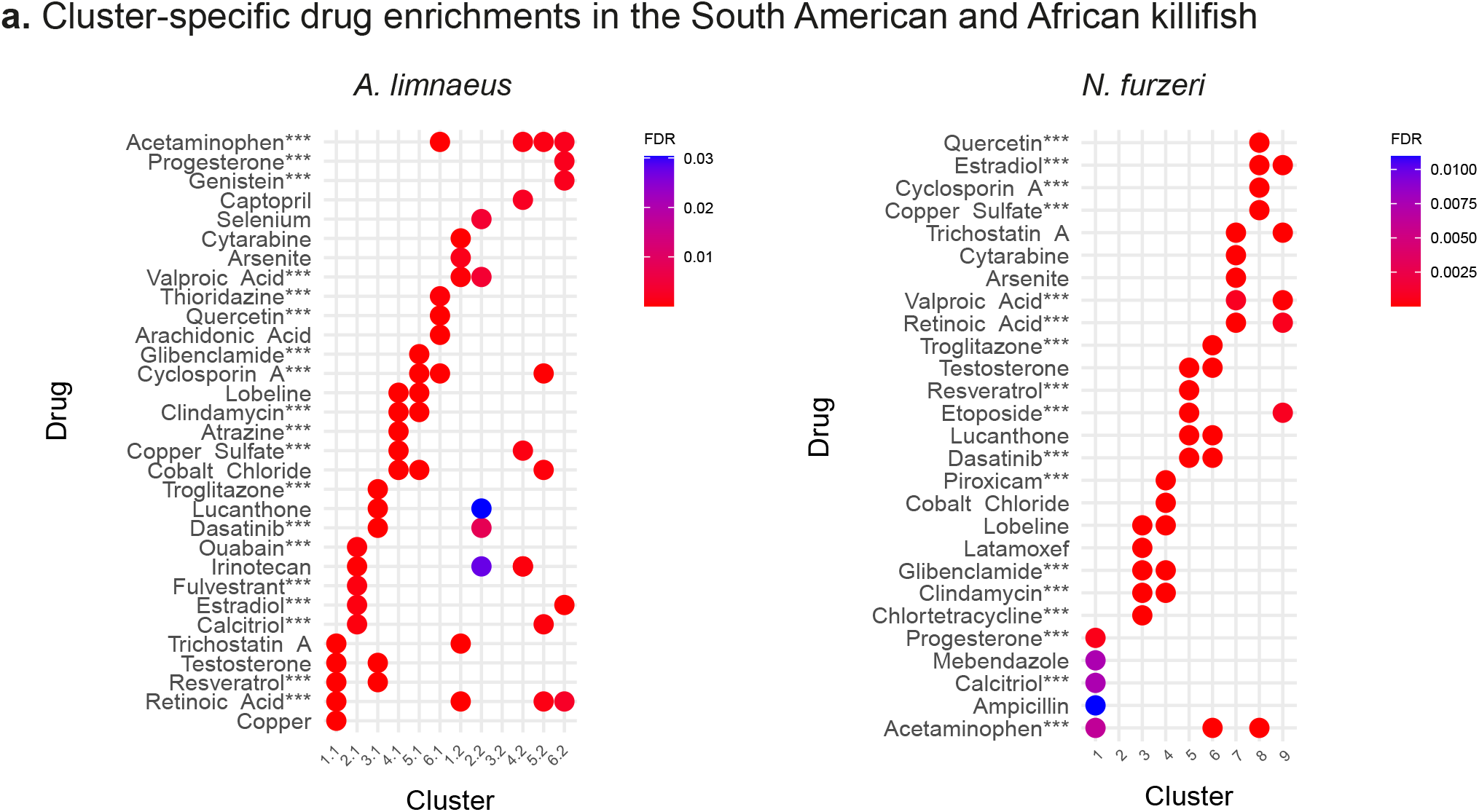
Cluster-specific drug enrichmentsa. Drug enrichments (rows) for each cluster (columns) in South American (left) and African (right) killifishes. Panels show the five most significant drugs per cluster, with 5% FDR-corrected p-values represented by color. Compounds selected for the final 111-compound functional screen are labelled with “***”.

**Figure S5.**
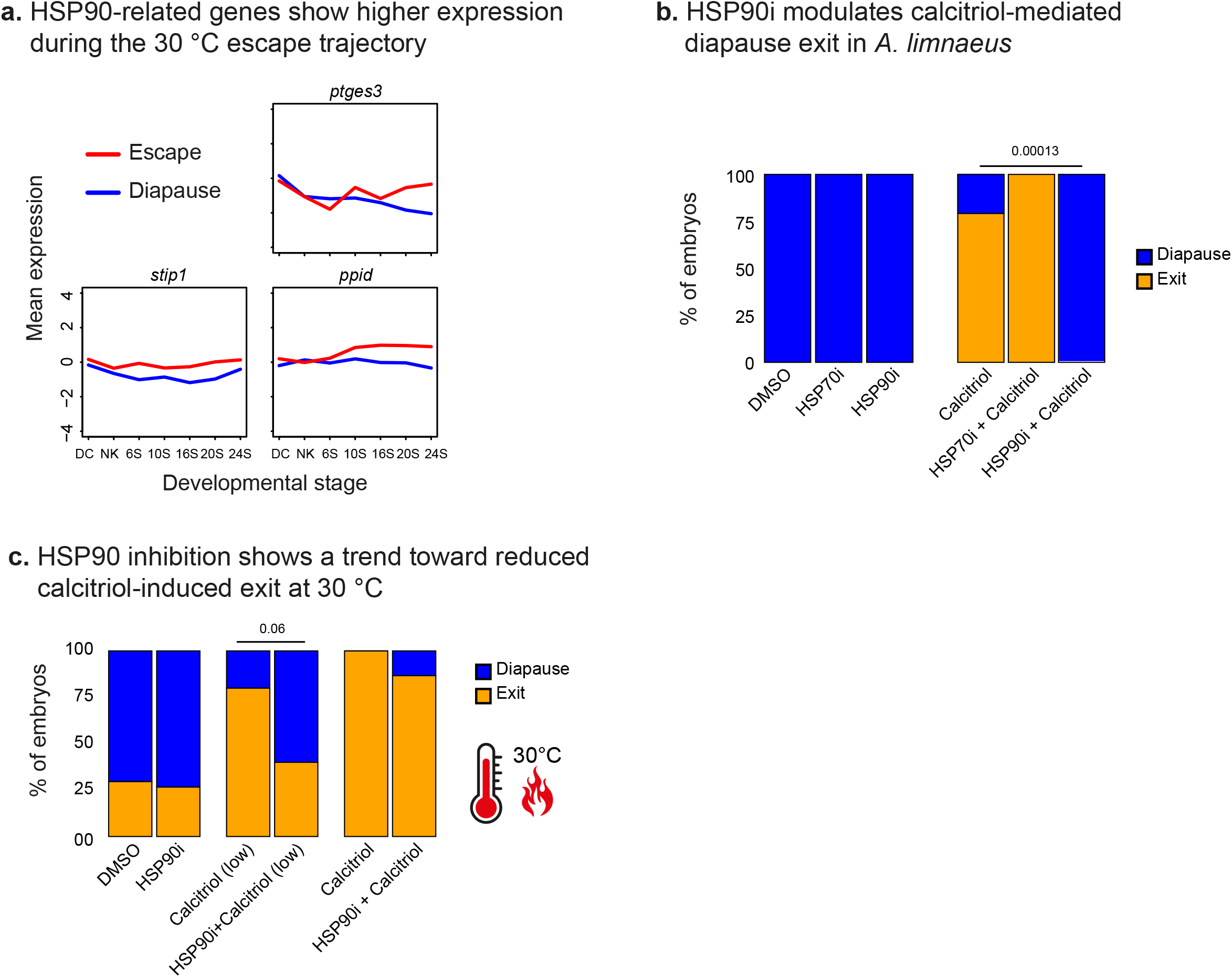
HSP90 activity modulates calcitriol-dependent diapause exit. **a.** Temporal expression profiles of four HSP90 co-factors from clusters showing higher expression at 30 °C compared to 20 °C (clusters in the top row of Fig. 3c). Lines show measured gene expression (y-axis) over time (x-axis) in 30 °C (red, escape) and 20 °C (blue, diapause) conditions. **b.** Pharmacological inhibition of HSP90 blocks calcitriol-dependent diapause exit in South American killifish. Bar graphs represent the proportion of embryos (y-axis) that either exit (orange) or maintain diapause (blue) following treatment as indicated at 25 °C after one week. P-values from Fisher’s exact tests are shown above. DMSO (2%), HSP70i (200µM), HSP90i (200µM), calcitriol (100nM), calcitriol (100nM) + HSP70i(200µM), calcitriol (100nM) + HSP90i(200µM). n=12 for all the bars (see **Methods**). **c.** African killifish embryos in diapause were incubated at 30 °C for one week under three treatments: DMSO control (n=27), 10 nM calcitriol (n=15), and 100 nM calcitriol (n=10). Each treatment was tested with or without 200 µM HSP90 inhibitor (n=15). Bar graphs represent the proportion of embryos (y-axis) that either exit (orange) or maintain diapause (blue) following treatment as indicated (x-axis) at 30 °C. P-values are shown above, calculated by a Fisher’s exact test.

**Supplementary Tables S1-16 Data relating to the entire manuscript**

**Movie S1 FUCCI embryos, calcitriol-dependent diapause exit**

This time-lapse record depicts two *N. furzeri* FUCCI embryos developing at 20°C for 72 hours, with a 1-hour imaging interval indicated in the top-left yellow timestamp. The embryos were imaged immediately following treatment with 200 nM Calcitriol after a 1-month incubation period at 20°C. Fluorescence is shown in two channels, where RFP indicates cells in the G0/G1 phases and GFP represents cells in the G2/M phases. A coordinated wave of proliferation, indicated by the GFP signal. This GFP burst is followed by further development beyond the diapause stage, characterized by pigmentation and black eyes.

## Methods

### African turquoise killifish strain, husbandry, and maintenance

The African turquoise killifish (GRZ strain) were housed as previously described^2,3,98,100,102,105,25,90–96^. Fish were housed at 28°C in a central filtration recirculating system with a 12 h light/dark cycle at the Hebrew University of Jerusalem (Aquazone ltd, Israel). Fish were exclusively fed with live Artemia until the age of 2 weeks (#109448, Primo), and starting week 3, fish were fed three times a day on weekdays (and once a day on weekends), with GEMMA Micro 500 Fish Diet (Skretting Zebrafish, USA), supplemented with Artemia once a day. In these conditions, killifish lifespan was approximately 4-6 months. All turquoise killifish care and use were approved by the Subcommittee on Research Animal Care at the Hebrew University of Jerusalem (IACUC protocol #NS-18-15397-2).

### Egg collection and induction of diapause

Mating couples were paired with sand trays for a week, and following collection, fertilized eggs were sterilized according to^97^. Embryos were incubated at a low temperature of 20 °C for 2 weeks to induce embryonic diapause, and their health was monitored twice a week. Embryos that escaped diapause during this period were removed. To synchronize embryos, we extended the incubation for one month^17,25^. For alternative developmental trajectories (**Figure S1a**), embryos were collected immediately following mating, and kept at 30 °C for 8-9 days, until the body axis was visible. For the normal-development trajectory, embryos remained at 30 °C; for the ‘rapid in-and-out’ trajectory, embryos were transferred to 20 °C for 2 days and then returned to 30 °C.

### *N. furzeri* Vitamin D₃ Treatment for FUCCI Imaging

Diapausing embryos of the *Nothobranchius furzeri* FUCCI transgenic line^24^ were used for live imaging experiments. Embryos approximately one-month post-fertilization (as described previously) were placed in groups of three per well in a U-bottom 96-well plate. Each well contained a total volume of 150 µL of Ringer’s solution (96724-100TAB, Sigma-Aldrich) supplemented with 200 nM Calcitriol (1α,25-dihydroxyvitamin D₃; CAS 32222-06-3; Sigma-Aldrich, Cat. No. 679101) in 2% DMSO. Images were acquired every 30 minutes in two fluorescent channels: mCherry and GFP. Imaging was conducted continuously over 48 hours using an Olympus IX83 microscope equipped with an Olympus DP28 camera. The ambient temperature was maintained at 20 °C and monitored throughout the experiment. All imaging experiments were performed across three independent batches of embryos on separate dates. For FUCCI imaging analysis, embryo layouts were manually cropped from the GFP channel, and intensity (I) was measured for each image. The intensity values were then normalized (I_norm_) to a range between 0 and 1 using:

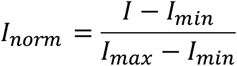

Image processing was performed in ImageJ, and illustrations as previously performed^108^.

### *N. furzeri* vitamin D treatment for RNA-seq

Diapausing *Nothobranchius furzeri* embryos (∼100 embryos, one-month post-fertilization) were distributed individually into wells of a U-bottom 96-well plate. Each embryo was treated with 200 nM Calcitriol (1α,25-dihydroxyvitamin D₃; CAS 32222-06-3; Sigma-Aldrich, Cat. No. 679101) dissolved in 2% DMSO. At 2.5-hour intervals, three embryos were collected and transferred into separate 1.5 mL microcentrifuge tubes containing 100 µL TRI Reagent (Sigma-Aldrich, Cat. No. T9424). Embryos were manually homogenized using sterile pestles until complete lysis. Tubes were immediately transferred to −80 °C and stored until RNA extraction.

### *N. furzeri* single-embryo RNA extraction

RNA extraction was performed following a modified version of the protocol described in^17^. Briefly, frozen, crushed *N. furzeri* embryos were thawed and further homogenized for 5 min at room temperature using zirconium oxide grinding beads and a TissueLyser LT (Qiagen). Samples were then incubated at room temperature for an additional 10 min and centrifuged for 15 min at 12,000 × g and 8 °C. The resulting supernatant (∼100 µL) was transferred to a clean 1.5 mL microcentrifuge tube, and 20 µL chloroform (CAS 67-66-3) was added. After vigorous mixing, the samples were centrifuged for 30 min at 21,000 × g and 8 °C to achieve phase separation. The upper aqueous phase was transferred to a new tube, and 25 µL isopropanol (CAS 67-63-0), 25 µL of salt precipitation solution (0.8 M sodium citrate, 1.2 M NaCl), and 1 µL GlycoBlue (Thermo Fisher Scientific, Cat. No. AM9516) were added. The mixture was vortexed and incubated overnight at −20 °C.

The next day, RNA was pelleted by centrifugation for 30 min at 21,000 × g and 8 °C. The supernatant was carefully discarded, and the pellet—visible as a small blue dot—was washed twice with 1 mL of 65% ethanol at room temperature. After each wash, tubes were gently inverted to dislodge the pellet and centrifuged for 30 min at 21,000 × g and 8 °C. Following the second wash, residual ethanol was removed, and the pellet was air-dried for 5 min at room temperature. RNA was dissolved in 30 µL RNase-free water (Thermo Fisher Scientific, CAS 7732-18-5) by incubation at 55 °C for 5 min, followed by gentle pipetting and brief centrifugation. RNA integrity was assessed using an Agilent Bioanalyzer, and samples were stored at −80 °C until use.

### RNA-Seq library preparation

RNA concentration and quality were determined by using an Agilent 2100 bioanalyser (Agilent Technologies)^103,109^. For each timepoint, the embryo with the best RNA quality was used for library preparation. Library preparation was performed using KAPA mRNA HyperPrep Kit (ROCHE-08105936001) according to the recommended protocols. Library concentrations were measured by Qubit (dsDNA HS, Q32854), and quality was measured by Tape Station (HS, 5067-5584). Libraries were sequenced by NextSeq 500 high output kit V2, 75-cycle single-end (Illumina, 20024906) using a NextSeq 500 machine (Illumina) with ∼30 million reads per sample.

### *N. furzeri* RNA sequencing analysis

Quality control and adapter trimming of the fastq sequence files were performed with FastQC (v0.11.8)^99^, fastx-toolkits (v0.0.13), Trim Galore! (v0.6.4)^101^, and Cutadapt (3.4)^104^. Options were set to remove Illumina TruSeq adapters and end sequences to retain high-quality bases with phred score > 20 and a remaining length > 20 bp. Successful processing was verified by re-running FastQC. Reads were mapped and quantified to the killifish genome Nfu_20140520^89,106^ using STAR 2.7.6a^107^. Gene expression was quantified by count per million (CPM) and log2 transformed.

### Temporal ordering of single-embryo RNA-Seq samples

We applied a PCA based approach to order the normalized single-embryo RNA-Seq samples by developmental time. Our approach is directly based on the BLIND approach^26^. Standardized log2 transformed normalized count per million (CPM) were used as input for principal component analysis (PCA, Matlab implementation). We considered the squared root of sum of squared differences as distance between every two samples on the principal components plane, and calculated the shortest path through the samples, which correspond to the developmental progress across the samples. Finding the shortest path on the principal components plane was done by solving the traveling salesman problem^110^, using the Matlab Problem-Based Optimization Workflow, with appropriate constraints. The resulting path is inferred as the developmental order of the samples.

Standardized log2 transformed normalized count per million (CPM) were used as input for principal component analysis of expression data (PCA, Matlab implementation). Gene set Enrichment analysis for PC1 was performed using GSEGO function from clusterProfiler R package. All the gene names in the RNAseq data are transformed to their human symbol and then ranked according to their weights on PC1 as calculated in the PCA analysis. Then the p-value is FDR corrected and filtered by corrected p-value <0.05 and ordered by their NES absolute value.

### *A. limnaeus* RNA sequencing data analysis

RNA-sequencing data from *Austrofundulus limnaeus* embryogenesis were downloaded from the National Center for Biotechnology Information (NCBI) Sequence Read Archive (SRA; https://www.ncbi.nlm.nih.gov/sra) under accession numbers SRX1032655, SRX1032657, SRX1032658, SRX1032664–SRX1032702 (BioProject ID PRJNA272154) (Romney et al., 2018). Raw sequencing data were converted to FASTQ format using the SRA Toolkit (v2.10.0, https://github.com/ncbi/sra-tools) with default parameters and trimmed using Trimmomatic (v0.39^111^) with the following settings: 2:30:7:1:true SLIDINGWINDOW:5:15 LEADING:20 TRAILING:20 MINLEN:25. Trimmed reads were aligned to the *A. limnaeus* reference genome (GCA_001266775.1^112^) using STAR (v2.7.5a^107^) with the parameters -- runThreadN 16 --outSAMtype BAM SortedByCoordinate. Mapped reads were quantified using HTSeq (v0.11.1^113^) via the htseq-count function with the options -f bam --order=pos -s yes --additional-attr=gene_name. For downstream analyses, the 7,000 most robustly expressed genes were selected based on total read counts. Normalization and transformation to log₂ counts per million (CPM) were performed using edgeR (v3.36.0^114^) with default parameters and the trimmed mean of M-values (TMM) normalization method. PCA for this data was performed as described above.

### RNA-Seq expression normalization

Temporal expression values (log2 transformed) were normalized by iteratively selecting a set of control genes, and normalizing all samples by their mean expression. Since standard RNA-seq normalization alone can be misleading for temporal samples, and particularly when overall mRNA amount in the embryo might change over time, we also normalized by control genes with known constant expression (very similar to qPCR normalization). Since we did not have a clear set of control genes, we applied an iterative approach^115^ to select those controls based on least varying genes, which has been benchmarked in previous studies^115–118^. Specifically, we fitted a linear regression model to log2-transformed means and standard deviations of genes, and selected genes with a standard deviation that is >2-fold lower than expected by the model. We repeated the normalization while number of selected controls increased. After normalization, only genes with the following criteria were selected for further analysis: log2(normalized FPKM)>2 in at least one temporal sample.

### Temporal function parameter fitting and selection

We used three alternative functions to describe temporal changes in gene expression levels (X) over time (t). A constant function (1 param):

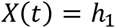

A single sigmoid function (4 param):

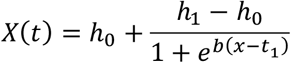

A double sigmoid “impulse” function (6 param):

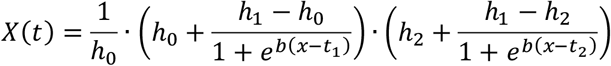

For each gene, each of 3 functions was fitted to temporal RNA-Seq data using non-linear least squares regression with multiple start values (Matlab implementation). A likelihood ratio test was applied to select optimal model, assuming gaussian additive noise with variance equals to the 70-th percentile variance of genes in the data.

### Selecting differential genes between two conditions

For each gene, we fit two alternative models: (1) “uniform” model fits one temporal function for all data (both conditions) as described above. Only the selected model is used. (2) “differential” model fits two temporal functions, one per condition as described above. We select between the two models using a likelihood ratio test to select between each gene, assuming gaussian additive noise with variance equals to the 70-th percentile variance of genes in the entire data (both conditions).

### Goodness-of-fit test

To assess the quality of each fit to the data, we use a Goodness of Fit test ^119^, assuming additive Gaussian noise with variance equal to the 70-th percentile variance of genes in the data. Degrees of freedom were specified by the number of measurements for the given condition, minus the number of parameters in the fitted function.

### Clustering of temporal expression data of *N. furzeri* RNA

We applied k-means clustering to the model-based expression predictions of genes, using the Python package *PyClustering*. This approach allowed us to smooth temporal patterns and eliminate sample-level noise, as previously suggested^120^. 5,230 genes were classified as temporally changing; 5,229 were assigned to the nine clusters. To cluster genes by expression pattern independently of expression magnitude, we used Pearson correlation as the distance metric. Model-predicted expression values were log₂-transformed, and fold change for each temporal sample of each gene (T_x_) was calculated relative to its expression value at the initial time point (T₀) as follows:

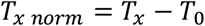

where T_x_ is the model-predicted log₂ expression at time point x, and T₀ is the model-predicted log₂ expression at the initial time point. The expression pattern of each cluster was defined as the mean fold-change profile of all genes within the cluster. The number of clusters (*k*) was first estimated by identifying the point at which adding more clusters led to only a slight improvement in within-cluster variance (the ‘elbow method’). Following this initial estimation, values of *k±2* were further tested to ensure that each cluster captured a distinct pattern of behavior.

### Clustering of *A. limnaeus* RNA sequencing data

We clustered the *A. limnaeus* RNA-seq data based on the dynamic behavior of each gene under two developmental trajectories: escape (30°C) and diapause (20 °C). For each gene in each condition, the fitted temporal expression model was used to classify its dynamic pattern. If the fitted model for a given condition was constant, the gene was labelled “constant” for that condition. For sigmoid-shaped models, the gene was classified as “up” if the model fitted an initial level lower than the second level, and “down” if the model fitted an initial level higher than the second level. For double-sigmoid models, classification was determined by the order of the minimum and maximum values: if the minimum occurred before the maximum, the gene was labelled “up”; if the maximum preceded the minimum, it was labelled “down.” This classification was performed independently for each condition, yielding nine possible combinations of dynamic behavior across hot and cold treatments: (up–up), (up–constant), (up–down), (constant–up), (constant–constant), (constant– down), (down–up), (down–constant), and (down–down). For the three symmetric combinations—up–up, constant–constant, and down–down—genes were further subdivided according to which condition showed higher overall expression. For each gene, we identified the time point with the maximum absolute difference in expression between the hot and cold conditions (Δ). Genes were then assigned to either the “escape-higher” or “diapause-higher” subclustered based on which condition exhibited higher expression at that time point.

### Functional enrichment analysis within clusters

Functional enrichment analysis was performed on gene clusters to identify significantly overrepresented biological pathways and processes. We used the *clusterProfiler* R package to conduct over-representation analysis (ORA) against curated gene sets from the Molecular Signatures Database (MSigDB), including KEGG, Reactome, Gene Ontology Biological Process, and Molecular Function categories. The background gene set consisted of all genes expressed in the RNA-seq samples, excluding those that did not fit the model. For each gene cluster of interest, enrichment was calculated with multiple testing correction using FDR method, with a false discovery rate cutoff of 0.05. GO terms were based on human GO annotations from mSigDB.

### Transcription factor binding site enrichment analysis within clusters

We used HOMER (Hypergeometric Optimization of Motif EnRichment, version 5.1, July 16, 2024) to identify enriched transcription factor binding motifs within each gene cluster. For each cluster, we analyzed a 1100 bp regulatory region surrounding the transcription start site (TSS)—specifically, 1000 bp upstream and 100 bp downstream—as annotated in the *Nothobranchius furzeri* genome assembly (Nfu_20140520). Enriched motifs were identified using HOMER’s findMotifsGenome.pl tool. The background set comprised the corresponding regulatory regions of all genes expressed in the RNA-seq dataset, excluding genes that failed model fitting. This motif enrichment analysis was performed independently for each cluster, as well as for combined clusters that showed similar functional enrichment or similar expression patterns.

### Comparative Cluster Overlap Analysis Using Permutation Testing

To assess the similarity between gene expression clusters derived from *A. limnaeus* and *N. furzeri*, we performed a permutation test evaluating the overlap of genes assigned to clusters in each species. First, we identified genes common to both datasets, restricted the analysis to these shared genes, and compiled their cluster assignments accordingly. For each pair of *A. limnaeus* and *N. furzeri* clusters, we calculated the observed number and ratio of overlapping genes. To evaluate the statistical significance of these overlaps, cluster labels in the *A. limnaeus* dataset were randomly shuffled 1,000 times, generating a null distribution of overlap counts for each cluster pair. Empirical p-values were calculated as the proportion of permuted overlaps greater than or equal to the observed overlap, and FDR multiple testing correction was applied. The overlap ratios calculated as ratio between the mutual genes in both clusters (intersection) and all the genes in both clusters (union), Jaccard index:

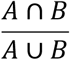

### Small molecules functional enrichment analysis within clusters

Drug-enrichment analysis was performed separately for each *N. furzeri* diapause-exit cluster and each *A. limnaeus* trajectory-specific cluster using the enrichR R package and the DSigDB database. The background gene set consisted of all expressed genes included in the RNA-seq analysis. For each cluster, enriched drug signatures were identified at FDR < 0.1 and compiled into a full candidate table (**Table S10**), yielding 440 compounds associated with the African killifish diapause-exit dataset and 496 compounds associated with the South American killifish diapause entry/escape dataset. Compound names were standardized and mapped to CAS numbers where possible. The two species-level lists were then merged, and compounds predicted in both species were prioritized as putative shared modulators of diapause-related state transitions. It is worth noting that a drug could be independently enriched in several clusters.

To assemble the experimental screening set, we applied additional filters, prioritizing compounds with stronger enrichment scores (FDR < 0.01), cross-species support, commercial availability in aliquoted format, and biological relevance to enriched pathways or candidate regulatory nodes. We excluded a subset of non-specific cytotoxic, proliferation-targeting chemotherapeutic, or poorly soluble/unstable compounds. This procedure produced the final set of 111 compounds used in the functional screen (**Tables S11–13**).

### Quantification of changes in biological age using BiT age

BiT Age was applied according to the detailed methods in ref.^87^, with adaptations for the *N. furzeri* and *A. limnaeus* transcriptomes. For the *N. furzeri*, the index for the alignment was created from Nothobranchius_furzeri.Nfu_20140520.cdna.all.fa, and with Nothobranchius_furzeri.Nfu_20140520.dna.toplevel.fa used as a decoy. For the *A. limnaeus* we used the samples from Romney et al^17^. To construct the clocks for the *N. furzeri* we used three pre-diapause samples from (Hu et al. 2020^21^) and for *A. limnaeus we used 3* samples at the DC stage maintained at 20°C^17^ to generate a stochastic noise-based time series. We confirmed that all clocks were comparable in trend, irrespective of the training sample, and provide a representative replicate in the paper. Raw counts were transformed using CPM-log10 (counts per million) and min-max scaled to define the ground state for stochastic noise simulation.

Briefly, for a given simulated age (*a*), we initialized the system at this ground state and applied a baseline noise. We then iteratively added age-related stochastic noise (N ∼ (0, 0.05)) for “*a”* iterations, with the gene expression values clipped to the [0, 1] interval after each addition to maintain the normalized range. The resulting simulated time series were binarized using a gene-specific median threshold. For each gene, expression values greater than the median across all simulated ages were set to 1, while values less than or equal to the median were set to 0. An ElasticNet model with the parameters random_state = 0, max_iter = 10000, l1_ratio = [0.1, 0.2, 0.3, 0.4, 0.5, 0.6, 0.7, 0.8, 0.9], alphas=[1] was then fitted to predict simulated age. Finally, the empirical diapause exit time-series data were log10-CPM transformed and binarized using the same median-based thresholding to serve as the input for the trained stochastic aging clocks. Because the clock is generated from a small ground-state reference set and models stochastic transcriptomic-noise accumulation, we interpret its output as predicted transcriptomic age/noise structure rather than calibrated organismal age.

### Transcriptomic staging using RAPToR

Transcriptomic staging was performed using the ‘RAPToR’ R package^41^. A dataset comprising seven developmental time points of *A. limnaeus,* reared at 30°C, was utilized as the reference to stage test samples incubated at 20°C^17^. The package was run using default parameters.

### Functional chemical screening in *A. limnaeus* and *N. furzeri* embryos

The final transcriptome-guided screening set consisted of 111 selected small molecules obtained from the Nancy and Stephen Grand Israel National Center for Personalized Medicine (G-INCPM), Rehovot, Israel. Candidate compounds were filtered before experimental testing to enrich for interpretable pathway perturbations and to reduce non-specific toxicity. Compounds were prioritized based on their pharmacological profiles, and a subset of highly cytotoxic, proliferation-targeting, or poorly water-soluble compounds was excluded. Compounds were dissolved in DMSO to a stock concentration of 100 µM and stored at −20 °C until use. Diapausing *Austrofundulus limnaeus* embryos (diapause II stage) were shipped from the USA, surface-sterilized with iodide, and maintained at 20 °C. For the activator screen, three embryos were placed per well in U-bottom 96-well plates containing 150 µL Ringer’s solution supplemented with 1 µM of the test compound and DMSO was adjusted to a final concentration of 2% in all wells (final concentration, without media change throughout the experiment). Plates were monitored weekly, and diapause exit was defined as the appearance of visible black eyes. For the inhibition screen, the procedure was identical except that each well additionally contained 100 nM Calcitriol (1α,25-dihydroxyvitamin D₃; CAS 32222-06-3; Sigma-Aldrich, Cat. No. 679101) along with the test compounds. The exit rate was calculated as the number of viable embryos that exited diapause divided by the total number of viable embryos per treatment condition after 3 weeks. Specifically, for the South American killifish, we tested 111 compounds using n = 9 embryos per compound (3 embryos per well across 3 wells) in the activation screen and n = 6 embryos per compound (3 embryos per well across 2 wells) in the inhibition screen. For the African killifish, we tested 31 compounds using n = 6 embryos per compound (3 embryos per well across 2 wells). Hit criteria were defined according to screening strategy rather than species. In the activator screen, compounds were classified as hits when ≥25% of viable embryos exited diapause after 3 weeks in the absence of calcitriol. In the inhibitor screen, compounds were classified as hits when they reduced calcitriol-induced diapause exit by ≥50% relative to the calcitriol-only control. The same inhibitor criterion was applied to the *N. furzeri* follow-up screen. Dead embryos were excluded from calculation of exit rates and from statistical analyses.

### Effect of HSP90 and HSP70 inhibitors on diapause exit

Ganetespib (HSP90 inhibitor; CAS 888216-25-9; BioTag) and VER-155008 (HSP70 inhibitor; CAS 1134156-31-2; BioTag) were prepared as 10 mM stocks in DMSO and stored at 4 °C. Embryos were assayed in U-bottom 96-well plates containing 150 μL Ringer’s solution, with inhibitors used at 200 μM and DMSO at a final concentration of 3%. For inhibition experiments, 100 nM calcitriol was added. Embryos were monitored for 2 weeks. Unless otherwise indicated in Table S14, n = 12 embryos were used per treatment see **Table S14**.

### Cells origin

Primary skin established from biopsies in the mid-1980s, were used in this study. Because their collection predated modern institutional ethics regulations, formal approval was not required at the time. These specific cell lines were previously characterized here^121^.

### Cell culture, transfection, and imaging

Cells (either human fibroblast or the murine NIH 3T3) were seeded on glass slides placed in 24-well plates and cultured for three days in DMEM (high glucose, with pyruvate; 41966029, Thermo Fisher Scientific) supplemented with charcoal-stripped fetal bovine serum (F6765-100ML, Sigma-Aldrich). Cells were then transfected with a VDR–RFP expression vector^72^ using Lipofectamine 3000 (L3000008, Thermo Fisher Scientific). For experimental conditions that included the HSP90 inhibitor Ganetespib (CAS: 888216-25-9, BioTag), it was added during transfection (10 nM). Twenty-four hours after transfection (efficiency ≈ 30%), cells were treated with 10 nM calcitriol (only for the experimental conditions that required calcitriol). Following an additional 8 hours of incubation, cells were fixed with 4% formaldehyde and stained with DAPI. Imaging was performed using an Olympus IX83 inverted microscope equipped with a DP28 digital camera.

### Convergence of relaxed and intensified selection across annual and non-annual killifish lineages

Gene-wise estimates of selection relaxation and intensification^85^ were compiled for two annual (*Callopanchax*, *Nothobranchius*) and two non-annual (*Aphyosemion*, *Scriptaphyosemion*) killifish genera. Comparative analyses between the two groups were performed by identifying overlaps between significant genes to detect opposing signatures of evolutionary pressure.

## Data availability

Newly generated *N. furzeri* raw and processed RNA-seq data are available in GEO under accession GSE307489. The published *A. limnaeus* RNA-seq data^17^ reanalyzed here are available in NCBI SRA under accessions SRX1032655, SRX1032657, SRX1032658, and SRX1032664–SRX1032702 (BioProject PRJNA272154). Source data are provided with this paper.

## Code availability

The code supporting the current study is available in the following GitHub repository: https://github.com/Harel-lab/Regulation-of-Diapause-exit, and the diapause temporal modeling is available at: https://github.com/rabanilab/Diapause_temporal_modeling

## Acknowledgements

We thank Yehu Moran, Sagiv Shifman, Yotam Drier, and the Harel and Rabani labs for stimulating discussion and feedback on the manuscript. We thank Fatma Idrees and Reem Barakat for help with killifish maintenance, Naomi Melamed-Book from the imaging facility (HUJI). Supported by ERC StG #101078188 (I.H.), the Zuckerman Program (I.H.), ISF 2178/19 (I.H.), Israeli Ministry of Science 3-17631 (I.H.), 3-16872 (I.H.), the Moore Foundation GBMF9341 (I.H.), BSF-NSF 2020611/2025832 (I.H./J.E.P.), the Israeli Ministry of Agriculture 12-16-0010 (I.H.), ISF 1176/21 (M.R.), ISF 746/25 (M.R.), ERC StG #852451 (M.R.), and NSF 2025832 (J.E.P.).

## Author Contributions

O.B.D., M.R., and I.H. designed the study. O.B.D., and I.R., performed experiments with help from E.M.D., and A.O.G., under the supervision of I.H. M.R. developed and applied the kinetic models. O.B.D. performed analysis of RNA sequencing with help from T.AT. and T.AR., and under the supervision of I.H. and M.R. E.M.D. performed the HSP90 experiments in the South American killifish and provided the South American killifish embryos for the small-molecule screen under the supervision of J.E.P. W.S., and D.H.M performed the BiT analysis under the supervision of B.S., M.T. performed drug validations in zebrafish under the supervision of M.R.. O.B.D., M.R, and I.H. wrote the manuscript. All authors commented on the manuscript.

